# Gliomas preferentially develop within the action-mode network

**DOI:** 10.64898/2026.01.05.697608

**Authors:** Weigang Cui, Jiayi Zhu, Zeya Yan, Jianxun Ren, Scott Marek, Veit M. Stoecklein, Tao Jiang, Hongbo Bao, Shengyu Fang, Sophia Stoecklein, Zhengting Cai, Wei Zhang, Xiaoxuan Fu, Evan M. Gordon, Danhong Wang, Yinyan Wang, Nico U.F. Dosenbach, Hesheng Liu

**Affiliations:** Changping Laboratory, Beijing, China; State Key Laboratory of Cognitive Neuroscience and Learning, Beijing Normal University, Beijing, China; Department of Neurosurgery, Beijing Tiantan Hospital, Capital Medical University, Beijing, China; Beijing Neurosurgical Institute, Capital Medical University, Beijing, China; Department of Psychiatry, Washington University School of Medicine, St Louis, MO, USA; Department of Neurosurgery, University Hospital, LMU Munich, Munich, Germany; Department of Radiology, University Hospital, LMU Munich, Munich, Germany; Integrative Neuroscience and Cognition Center, Université Paris Cité, Paris, France; Mallinckrodt Institute of Radiology, Washington University School of Medicine, St Louis, MO, USA; Allied Labs for Imaging Guided Neurotherapies (ALIGN), Washington University School of Medicine, St Louis, MO, USA; Department of Neurology, Washington University School of Medicine, St Louis, MO, USA; Department of Biomedical Engineering, Washington University in St. Louis, St Louis, MO, USA; Department of Psychological and Brain Sciences, Washington University in St. Louis, St Louis, MO, USA; Department of Pediatrics, Washington University School of Medicine, St Louis, MO, USA; Biomedical Pioneering Innovation Center, Peking University, Beijing, China

## Abstract

Gliomas tend to arise in specific brain regions and may integrate into functional circuits, suggesting they could be regulated by brain activity. However, it remains unclear whether glioma growth is related to system-level brain networks. Analyzing neuroimaging data from three datasets including 1,310 patients with cerebral gliomas, we identified and replicated a functionally connected glioma network, which overlaps with the action-mode network (AMN), somatomotor network (SMN), and action-related subcortical regions. Resting-state functional connectivity (RSFC) of the AMN successfully predicted the location of glioma occurrence in two independent datasets with complex tumor distributions. Remarkably, no patient had a glioma entirely outside the AMN, and over 89% of patients exhibited gliomas with at least 50% overlap with the network. Moreover, the spatial overlap between glioma location and the AMN demonstrated significant prognostic value in survival analyses, with higher AMN-tumor overlap associated with poorer overall survival. Notably, the acetylcholine transporter, a key player in glioma pathogenesis that drives transcriptional reprogramming, showed an expression pattern overlapping with the AMN. Meta-analytic annotations further linked the glioma network to processes of action initiation, execution, and feedback. These findings indicate that gliomas preferentially arise in circuits involved in action and highlight the central role of the AMN in glioma pathophysiology and growth.

---

Gliomas are characterized by rapid cellular proliferation and extensive infiltration into surrounding brain tissues, making them highly aggressive and challenging to treat^1–4^. Unlike most tumors that predominantly affect older adults^5,6^, gliomas lack established cancer risk factors such as environmental carcinogens, chronic inflammatory conditions and heredity, with lower-grade subtypes targeting young to middle-aged adults^7,8^, suggesting unique oncogenic drivers embedded within the brain^9,10^. Recently, gliomas have been increasingly recognized as dynamic entities influenced by bidirectional interactions with their neural microenvironment^11–17^. Emerging evidence from electrocorticography (ECoG) and task-evoked functional magnetic resonance imaging (fMRI) has revealed intratumoral neural responses evoked by goal-directed behaviors, such as speech production^18,19^ and movement execution^20^. These findings highlight that glioma can integrate into functional circuits and thus may be influenced by common task-related processes. Neuron-glioma interactions, mediated by paracrine signaling and neurotransmitter-dependent (e.g., glutamate and acetylcholine) synapses^21–27^, contribute to neuronal hyperexcitability within tumors^11–17,27^. This hyperactivity has been associated with accelerated tumor growth and proliferation^18,23,27,28^. Collectively, these insights suggest that glioma distribution may, in part, be shaped by neuronal activity across different brain regions.

Goal-directed activity is associated with increased arousal, action planning, and feedback integration^29–31^. An action-related circuit, called the action-mode network (AMN), supports various goal-directed behavior^32^. This network spans the cerebral cortex, subcortex, and cerebellum, with key nodes located in the dorsal anterior cingulate cortex (dACC), anterior insula, supplementary motor area (SMA), supramarginal gyrus (SMG), inferior frontal gyrus (IFG), anterior putamen and ventral intermediate thalamus^29,32–34^. Task-evoked fMRI studies consistently highlight AMN’s involvement in goal-directed activities such as executive control^29,35–37^, motor control^35,38^, and pain monitoring^39,40^. Additionally, AMN is strongly connected to the somatomotor network (SMN), particularly within its recently defined somato-cognitive action network (SCAN)^41^. Thus, the AMN, responsible for action initiation and planning, and the SMN, responsible for execution, form a highly integrated circuit that is consistently engaged during goal-directed tasks.

Gliomas exhibit a non-random spatial distribution, with a predilection for highly connected brain regions such as the insula, temporal cortex, and putamen^42–44^. Emerging evidence indicates that these tumors preferentially localize to functional hubs — regions characterized by high connectivity and centrality^3,4,45^. This spatial bias is further supported by magnetoencephalography (MEG) studies, which associate glioma occurrence with regions of elevated broadband power^43^. Nevertheless, the relationship between glioma-prone regions and functional brain networks has not been established.

To investigate the relationship between glioma growth and functional brain organization, we developed an approach called ‘glioma network mapping’, which is adapted from lesion network mapping^46–50^. This method offers a framework to reconcile heterogeneous glioma occurrence patterns and provides key insights into their network-based localization preferences. Using this approach, we mapped the glioma network across three independent neuroimaging datasets comprising 1,310 patients with cerebral gliomas. We investigated whether these glioma networks converged onto the AMN and used the AMN to predict glioma occurrence in new datasets featuring complex cases (*n* = 87, including cerebellar and multifocal gliomas). Additionally, we assessed survival outcomes (*n* = 285 across two datasets) based on the overlap between glioma networks and the AMN. Finally, we characterized the glioma network by neurotransmitter profiling^51,52^ and comparative functional pattern analytics^53^, to elucidate the neurochemical and functional properties of the identified glioma network.

## Gliomas are spatially distributed but functionally connected

We analyzed the spatial distribution of glioma locations in the cerebral cortex across three independent datasets (Tiantan: *n* = 830; BraTS: *n* = 345; UPenn: *n* = 135; see Supplementary Table 1 for detailed demographic and clinical information), revealing heterogeneous anatomical locations of gliomas (Fig. 1a). Gliomas were rare in the occipital cortex but occurred more frequently in the insula and temporal cortex (Fig. 1b). Nevertheless, anatomical overlap of the glioma locations across individuals, after registering to a common atlas (calculated as the percentage of patients showing a glioma at a given brain voxel) was quite limited, with the highest overlaps being 14.5% for Tiantan dataset, 13.0% for BraTS dataset, and 17.0 % for UPenn dataset (Supplementary Table 2).

**Fig. 1.**
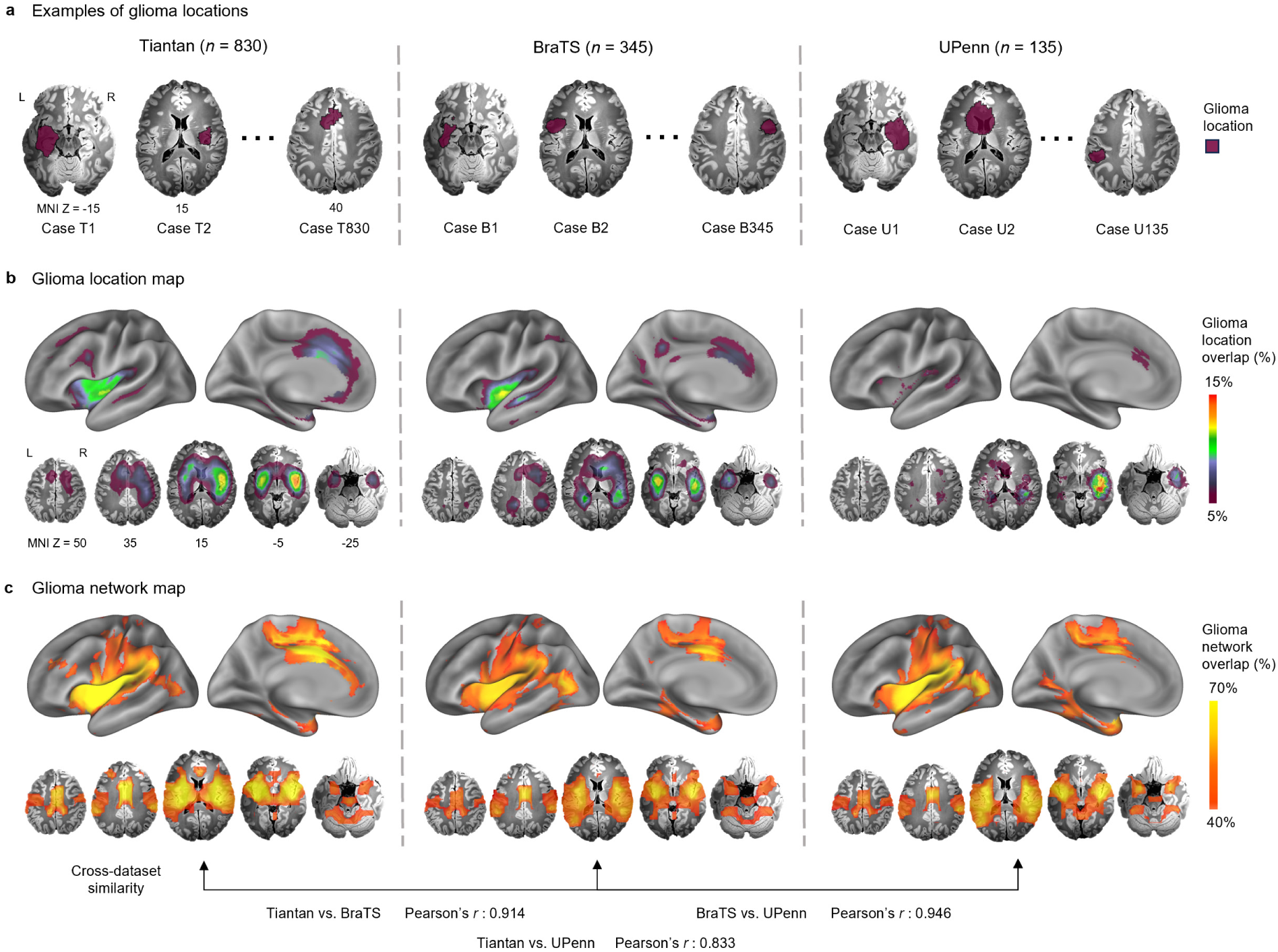
Anatomical distribution and functional connectivity pattern of gliomas. **a,** Examples of glioma anatomical locations (purple) are shown on a standardized Montreal Neurological Institute (MNI) brain atlas. **b,** Glioma anatomical location maps are displayed on brain surfaces (top, see Extended Data Fig. 2 for whole-brain surface map) and in volumetric space (bottom) for Tiantan (*n* = 830), BraTS (*n* = 345) and UPenn (*n* = 135) datasets, respectively. The color scale indicates glioma anatomical overlap across patients, representing the percentage of patients showing a glioma at each voxel. **c,** Glioma network maps, derived from normative resting-state functional connectivity (RSFC) of glioma locations (see “glioma network mapping” in Methods), are displayed for the three datasets (see Extended Data Fig. 2 for whole-brain surface map), with Pearson’s correlation coefficients between the maps shown in the bottom (all one-sided *P*_spin_ < 0.001). The color scale reflects the overlap of glioma-connected networks across patients, calculated as the percentage of patients exhibiting significant functional connectivity between the glioma and a given voxel.

Recent progress in lesion network mapping has revealed that brain lesions can occur in an anatomically distributed, but functionally connected network and lead to the same neuropsychiatric symptom^46–50^. We thus applied a glioma network mapping approach (see Methods and Extended Data Fig. 1 for algorithm framework) to explore the functional relationship of those anatomically distributed cerebral gliomas (Fig. 1c). The analyses revealed a functionally coherent glioma network which included the bilateral insula, posterior inferior frontal gyrus, posterior middle frontal gyrus, posterior and dorsal anterior cingulate gyri, supramarginal gyrus, anterior prefrontal cortex, and subcortical structures such as the putamen, globus pallidus, and thalamus. We quantified cross-individual functional overlap as the percentage of patients exhibiting significant positive connectivity with the glioma at a given brain voxel. The right insula was most consistently found as the functional focus of the glioma network, with this functional overlap of 76.4% in the Tiantan dataset, 70.1% in the BraTS dataset, and 71.1% in the UPenn dataset (see Supplementary Table 3 for foci coordinates). Negative connectivity with the glioma locations was mainly observed in the lateral occipital cortex, parietal lobe, inferior temporal sulcus, and precuneus (Supplementary Fig. 1). The glioma network map exhibited strong consistency across datasets, with high spatial correlation between maps (Fig. 1c; Tiantan vs. BraTS: Pearson’s *r* = 0.914, CI = [0.888, 0.934]; Tiantan vs. UPenn: Pearson’s *r* = 0.833, CI = [0.786, 0.870]; BraTS vs. UPenn: Pearson’s *r* = 0.946, CI = [0.930, 0.959]). These correlations were significantly greater than expected by chance (spin test, 10,000 permutations; all one-sided *P*_spin_ < 0.001, false discovery rate (FDR) corrected), indicating a shared functional pattern in glioma growth preference across ethnically diverse patient populations.

## The glioma network overlaps with the action-mode network

Given the high cross-dataset consistency of the glioma network, we constructed a comprehensive, cortico-subcortical version by aggregating data from all 1,310 patients across the three datasets (Fig. 2a-c). The spatial topography of the overall glioma network was highly stable across a range of statistical thresholds used to define functionally-connected regions (i.e., |*t*| > 5, 7, and 9), with strong spatial correlations between the resulting network maps (all Pearson’s *r* > 0.950, *P* < 0.001, FDR-corrected; Extended Data Fig. 3). In the cerebral cortex, the glioma network was mainly localized to the dorsal anterior cingulate cortex, anterior insula, and supramarginal gyrus (Fig. 2a; see Supplementary Table 4 for foci coordinates). In the cerebellum, the network encompassed an anterior portion (lobes IV, V, and VI) and a posterior portion (lobes VIIIA and VIIIB; Fig. 2b and Supplementary Table 4). Within the subcortical nuclei, the anterior putamen, globus pallidus and dorsal thalamus exhibited strongest glioma connectivity (Fig. 2c and Supplementary Table 4).

**Fig. 2.**
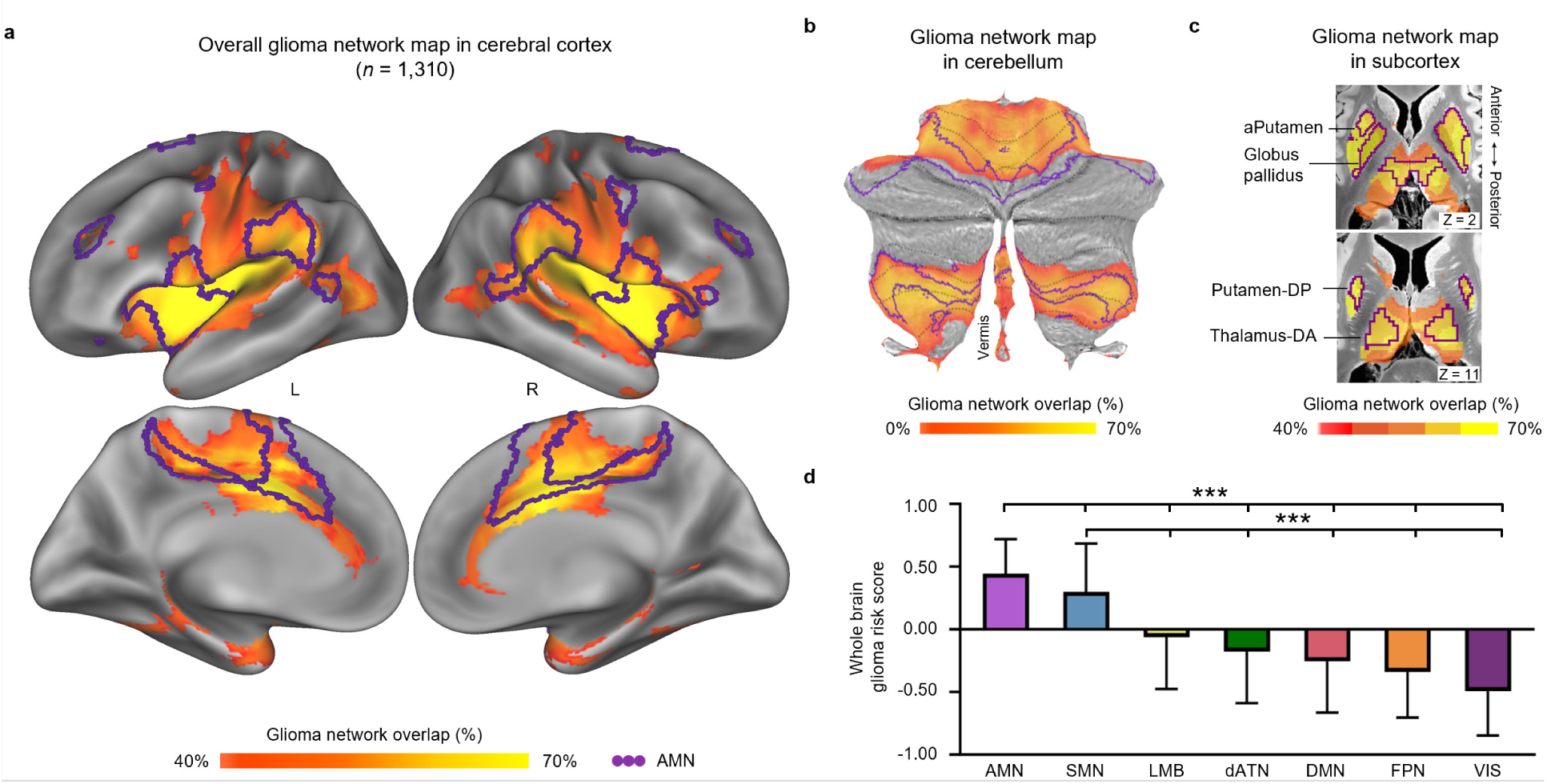
The cortico-subcortical distribution of glioma connectivity and risk score. **a,** Glioma network in the cerebral cortex based on all 1,310 patients from the Tiantan, BraTS and UPenn datasets (see Supplementary Fig.2 for the cortical and subcortical maps with different thresholds). The color scale reflects the overlap of glioma-connected networks across patients, calculated as the percentage of patients exhibiting significant connectivity between the glioma and a given voxel. **b** Glioma network in cerebellar regions (flat map). **c** Glioma network in subcortical regions. **d,** Glioma risk scores averaged in seven canonical networks across the whole brain. The AMN (action-mode network) exhibits the highest glioma risk score, followed by the SMN (somatomotor network; two-tailed independent *t*-tests with AMN or SMN, *** *P* < 0.001, FDR-corrected; see Extended Data Fig. 4 for detailed atlas^54–57^). In **a**-**c**, the AMN is outlined in purple. Bars indicate mean and error bars indicate standard deviation, s.d. a: anterior. DA: dorsoanterior. DP: dorsoposterior.

To further assess the functional relevance of the glioma network, we computed a glioma risk score for each brain voxel by measuring cross-individual overlap in the glioma-connected network, then evaluated this score within seven canonical networks from a standardized functional atlas (see Methods and Extended Data Fig. 4 for detailed atlas and scores across the cortex, cerebellum, and subcortex)^54–56^. The AMN exhibited the highest glioma risk score (0.445 ± 0.276), followed by the SMN (0.299 ± 0.385), which demonstrated significantly greater overlap with the glioma network compared to other functional networks (Fig. 2d; two-tailed independent *t*-test, all *P* < 0.001, FDR-corrected). Within the SMN, the SCAN exhibited a glioma risk score nearly twofold higher (0.251 ± 0.344) compared to the motor effector network (including foot, hand, and mouth regions; 0.146 ± 0.411; Extended Data Fig. 5). The visual network demonstrated the lowest glioma risk (-0.493 ± 0.354; see Supplementary Table 5 for glioma risk scores in other networks).

To account for potential variability across different tumor subtypes, we replicated the glioma network mapping analysis in two subgroups, patients with low- versus high-grade gliomas (low-grade: *n* = 432, age = 38.9 ± 12.2 years; high-grade: *n* = 398, age = 49.5 ± 15.2 years), and those with isocitrate dehydrogenase (IDH)-mutant versus wild-type gliomas (IDH-mutant: *n* = 477, age = 41.7 ± 12.6 years; IDH-wild: *n* = 353, age = 47.0 ± 16.7 years). While some inter-group differences were observed, the affected regions consistently overlapped with the AMN (Extended Data Fig. 6 and Supplementary Table 5).

## The AMN predicts the probability of glioma occurrence in new datasets

Building on the strong spatial correspondence between the glioma network and the AMN across the cortex, subcortex, and cerebellum, we further investigated whether the AMN could predict the location of gliomas in new datasets. We analyzed two independent datasets with more complex glioma distributions: multifocal gliomas (*n* = 38, age = 52.4 ± 14.9 years) and cerebellar gliomas (*n* = 49, age = 43.2 ± 17.2 years; Supplementary Table 1). The normative resting-state functional connectivity (RSFC) of the AMN was used as a predictive indicator (see Methods). Strikingly, in both datasets, all gliomas exhibited strong spatial overlap with the AMN (Fig. 3a, c). For multifocal gliomas, 89.5% of patients (34/38) showed tumors overlapping with >50% of AMN regions, with nearly one-third (31.6%, 12/38) demonstrating >75% overlap (Fig. 3b). Similarly, cerebellar gliomas displayed even stronger alignment: 91.8% of patients (45/49) had >50% tumor-AMN overlap, and nearly half (49.0%, 24/49) exceeded 75% overlap (Fig. 3d).

**Fig. 3.**
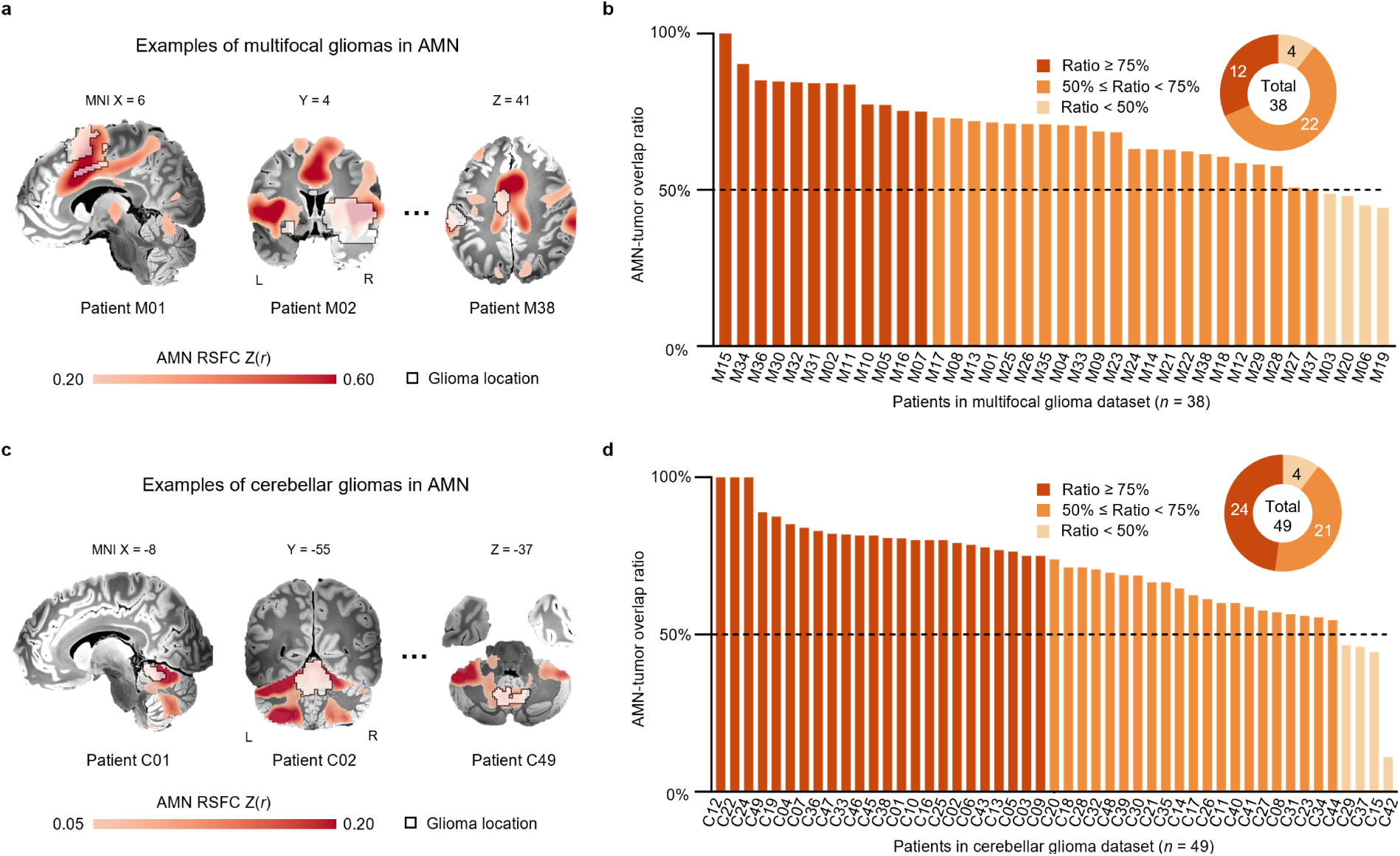
Prediction of glioma occurrence in independent datasets with complex tumor distributions. **a,** Examples of multifocal gliomas (black outline) overlaid on the AMN-connected regions. Functional connectivity with the AMN is shown in the brain volume. **b,** The AMN predicts multifocal glioma occurrence (*n* = 38, age = 52.1 ± 13.0 years), with 34 patients (89.5%) exhibiting more than 50% AMN-tumor overlap, and 12 patients (31.6%) surpassing 75% overlap. **c,** Examples of cerebellar gliomas (black outline) overlaid on the AMN functional connectivity regions, masked by the cerebellar region. **d,** The AMN predicts cerebellar glioma occurrence (*n* = 49, age = 43.2 ± 17.2 years), with 45 patients (91.8%) having more than 50% of their tumor volume within AMN regions, and 24 patients (49.0%) showing over 75% overlap.

To investigate the specificity of the prediction, we compared the performance of the AMN with that of six canonical functional networks^54^. Across both datasets, tumor locations exhibited significantly stronger alignment with the AMN than with any other network (two-tailed paired *t*-test, all *P* < 0.001, FDR-corrected; Extended Data Fig. 7a, b). To further determine whether these spatial correspondences beyond chance levels, we performed a glioma-null permutation test (1,000 permutations; see Methods and Extended Data Fig. 7c). In both datasets, observed AMN-tumor overlap ratios significantly exceeded those derived from randomly positioned, volume-matched control regions of interest (ROIs). For multifocal gliomas, the mean overlap with AMN was 68.9% ± 13.2%, compared to 47.4% ± 1.9% under the null model (*P*_perm._ < 0.001; Extended Data Fig. 7d). In the cerebellar dataset, the observed AMN-tumor overlap was 71.0% ± 6.0%, relative to 50.4% ± 4.0% in the null distribution (*P*_perm._ = 0.006; Extended Data Fig. 7e). These findings support that AMN is a robust predictor of glioma occurrence, even in anatomically complex cases.

## Overlap between glioma and the AMN predicts survival times

We hypothesized that gliomas with greater involvement of the AMN would be associated with poorer clinical outcomes, thus AMN-tumor overlap may serve as a prognostic marker for patient survival. To test this, glioma patients with available survival data from two independent datasets (Tiantan and UPenn; see Supplementary Table 6 for demographics) were stratified into high- and low-overlap groups based on the AMN-FC-weighted overlap score (see Methods). The cutoff threshold for stratification was independently determined in each dataset using maximally selected rank statistics^58^ (see Supplementary Fig. 3 and Methods for details). Kaplan-Meier survival analysis demonstrated that, across both datasets, high-overlap patients had significantly shorter overall survival than low-overlap patients (Fig. 4a, c). In the Tiantan dataset (*n* = 164, age = 45.4 ± 13.1 years), high-overlap patients (*n* = 75, age = 46.0 ± 13.6 years; AMN-FC-weighted overlap score: 0.160 ± 0.041) had a median overall survival time of 28.50 months, markedly shorter than the 50.34 months observed in the low-overlap patients (*n* = 89, age = 44.9 ± 12.7 years; AMN-FC-weighted overlap score: 0.066 ± 0.026; log-rank test for Kaplan-Meier survival analysis, *P* = 0.0010; Fig. 4a). Univariate Cox regression further confirmed the prognostic significance of the AMN-tumor overlap (hazard ratio (HR) = 1.906, *P* = 0.0012, confidence interval (CI) = [1.290, 2.815]). In addition, age (HR = 1.022, *P* = 0.0061, CI = [1.006, 1.037]) was also a significant predictor of survival (Error band represent 95% CI, Fig. 4b; see Supplementary Table 7 for details on all variables). After adjusting for potential confounders including age, gender, glioma grade and tumor volume, multivariate Cox regression identified the AMN-FC-weighted overlap score as a strong predictor of overall survival (HR = 1.829, *P* = 0.0025, CI = [1.236, 2.707]; Supplementary Table 7).

**Fig. 4.**
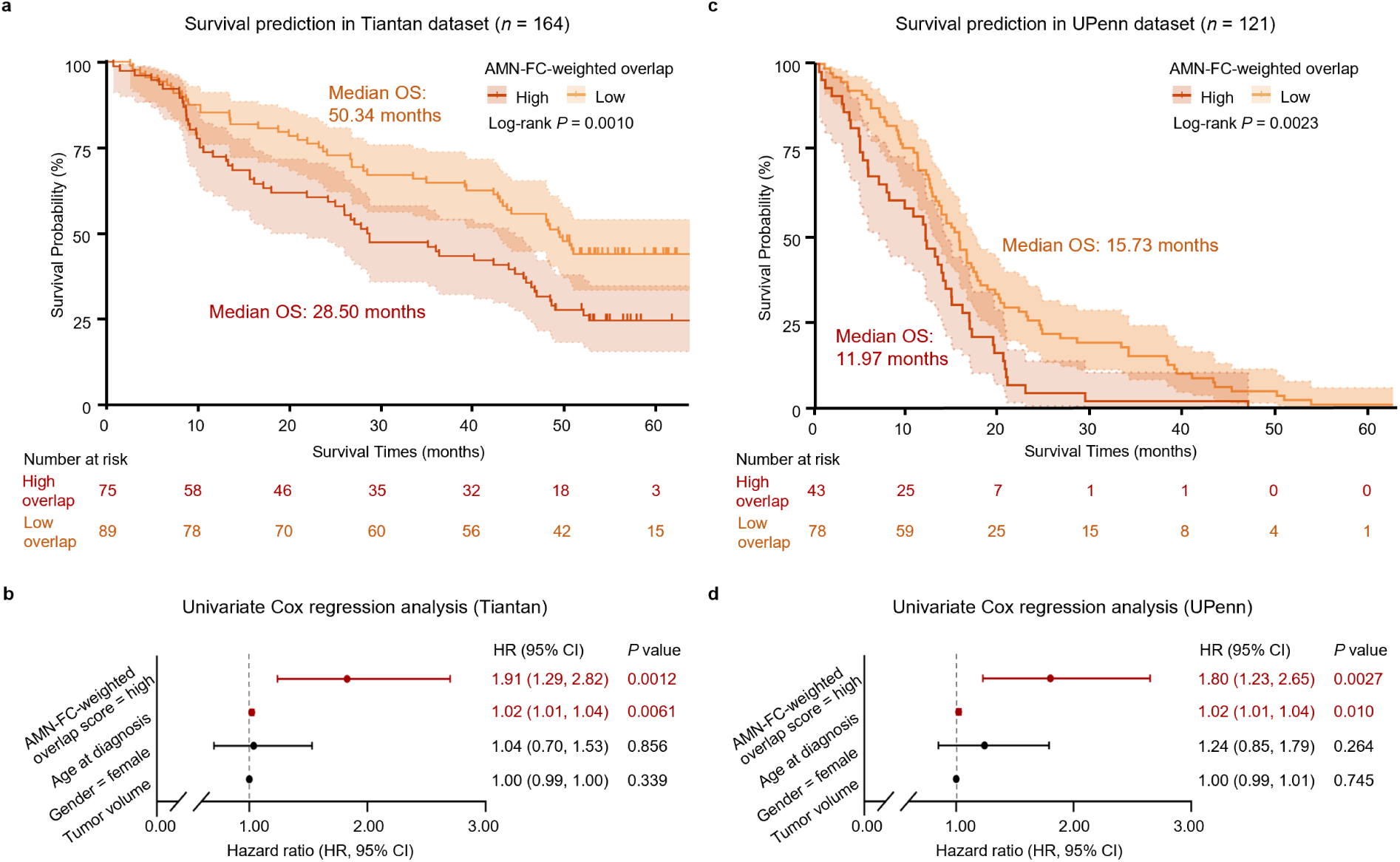
Prediction of patient survival time by AMN-FC-weighted overlap score. **a,** Kaplan-Meier survival curves comparing overall survival between high and low AMN-FC-weighted groups in the Tiantan dataset (*n* = 164, age = 45.4 ± 13.1 years). High-overlap patients (red; *n* = 75, age = 46.0 ± 13.6 years; AMN-FC-weighted overlap score: 0.160 ± 0.041; median survival time: 28.50 months) exhibited significantly shorter survival compared to low-overlap patients (orange; *n* = 89, age = 44.9 ± 12.7 years; AMN-FC-weighted overlap score: 0.066 ± 0.026; median survival time: 50.34 months) (log-rank test, *P* = 0.0010). Vertical ticks indicate censored events; shaded error bands represent 95% confidence intervals (CIs). **b,** Forest plot showing results from univariate Cox regression analysis in the Tiantan dataset. A higher AMN-FC-weighted overlap score demonstrated a significantly elevated hazard of death (HR = 1.91, *P* = 0.0012, CI = [1.29, 2.82]) relative to the low-overlap group. Age was also a significant predictor (HR = 1.02, *P* = 0.0061, CI = [1.01, 1.04]). Gender and tumor volume were not significantly associated with survival. Hazard ratios are indicated by filled circles, and whiskers represent 95% CIs. **c,** Kaplan-Meier survival curves comparing overall survival in the UPenn dataset (*n* = 121, age = 60.5 ± 10.5 years). High-overlap patients (red; *n* = 43, age = 61.6 ± 9.20 years; AMN-FC-weighted overlap score: 0.168 ± 0.061; median survival time: 11.97 months) exhibited significantly shorter overall survival than low-overlap patients (orange; *n* = 78, age = 59.9 ± 11.7 years; AMN-FC-weighted overlap score: 0.043 ± 0.033; median survival time: 15.73 months) (log-rank test, *P* = 0.0023). **d,** Forest plot showing results from univariate Cox regression analysis in the UPenn dataset. A high AMN-FC-weighted overlap score was associated with a significantly increased hazard of death (HR = 1.80, *P* = 0.0027, CI = [1.23, 2.65]). Age was also a significant predictor (HR = 1.02, *P* = 0.010, CI = [1.01, 1.04]). Gender and tumor volume did not show an association with survival time. Full regression results are provided in Supplementary Table 7.

The UPenn dataset (*n* = 121, age = 60.5 ± 10.5 years) comprised an older patient cohort with a higher prevalence of glioblastomas^59^. Kaplan-Meier analysis similarly revealed that the high-overlap patients (*n* = 43, age = 61.6 ± 9.20 years; AMN-FC-weighted overlap score: 0.168 ± 0.061) had a median survival of 11.97 months, significantly shorter than 15.73 months in the low-overlap patients (*n* = 78, age = 59.9 ± 11.7 years; AMN-FC-weighted overlap score: 0.043 ± 0.033; log-rank test for Kaplan-Meier survival analysis, *P* = 0.0023; Fig. 4c). Univariate Cox regression identified both the AMN-FC-weighted overlap score (HR = 1.803, *P* = 0.0027, CI = [1.227, 2.650]) and age (HR = 1.022, *P* = 0.010, CI = [1.005, 1.039]) as significant predictors of survival (Fig. 4d). After controlling for age, gender, and tumor volume, multivariate analysis showed that AMN-FC-weighted overlap score remained a highly significant predictor (HR = 1.798, *P* = 0.0030, CI = [1.220, 2.650]; Supplementary Table 7).

## Acetylcholine transporter is dense within the glioma network

Emerging evidence from calcium imaging and transcriptomic analyses indicate that acetylcholine (ACh) modulates glioma cell migration and tumor progression via the M3 metabotropic receptor (CHRM3)^21–24,59^. We obtained surface-based maps for 18 neurotransmitter receptors and transporters from the Neuromap toolbox^51,52^ and assessed their correlations with the glioma network. The vesicular acetylcholine transporter (VAChT), which is predominantly localized within the AMN (purple-highlighted dots in Fig. 5a) and plays a crucial role in the release of ACh at synaptic terminals, exhibited the highest positive correlation (Fig. 5a; Pearson’s *r* = 0.652, one-sided *P*_spin_ = 0.009, CI = [0.564, 0.722], spin test with 10,000 permutations, FDR-corrected). The glioma network showed significant spatial correlation with the densities of two additional neurotransmitter transporters (one-sided *P*_spin_ < 0.05, FDR-corrected; Fig. 5b), including serotonin transporter 5-HTT (Pearson’s *r* = 0.498, *P*_spin_ = 0.041, CI = [0.390, 0.593]) and dopamine transporter DAT (Pearson’s *r* = 0.472, *P*_spin_ = 0.048, CI = [0.362, 0.571]). In contrast, the GABA receptor (GABA_A/BZ_) exhibited negative spatial correlation with the glioma network (Pearson’s *r* = -0.317, *P*_spin_ = 0.238, CI = [-0.191, -0.433]).

**Fig. 5.**
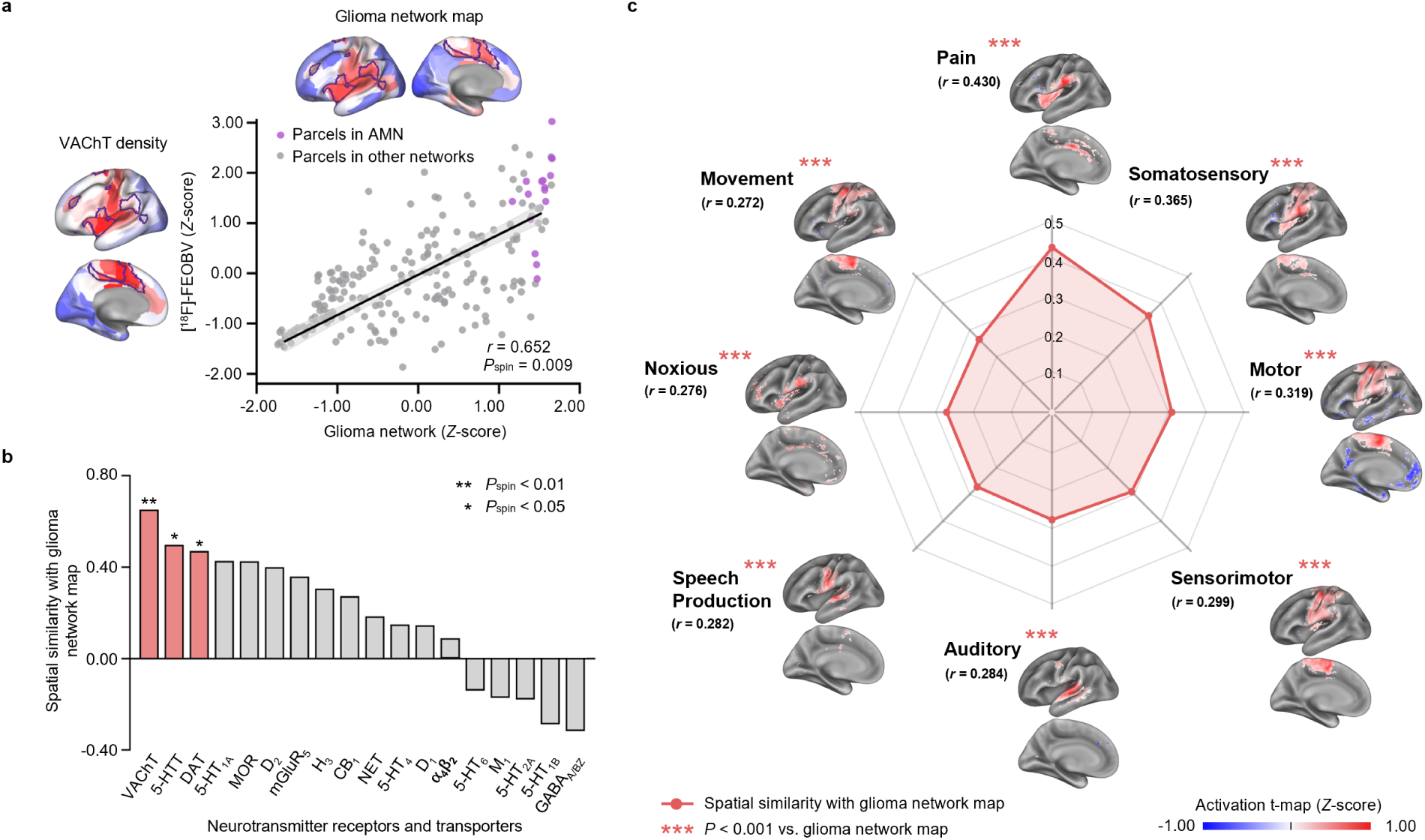
Neurochemical and functional signatures of the glioma network. **a,** Cortical surface visualization of the glioma network map and the distribution of VAChT density derived from PET imaging using the radiotracer ^18^F-fluoroethoxybenzovesamicol (FEOBV; 213 parcels, see Supplementary Fig.5 for details). The glioma network shows a significant spatial correlation with VAChT density (Pearson’s *r* = 0.652, one-sided *P*_spin_ = 0.009, CI = [0.564, 0.722], spin test with 10,000 permutations). Surface parcels belonging to the AMN are highlighted as purple dots in the scatter plot. **b,** Bar plot illustrates the spatial similarity between the glioma network and density maps of 18 neurotransmitter receptors/transporters. The top three maps with significant correlations are highlighted in red (** *P*_spin_ < 0.01, * *P*_spin_ < 0.05, FDR-corrected; one-sided), including transporters related to acetylcholine, serotonin and dopamine. **c,** Spatial similarity between the glioma network and the top eight behavior-related functional activation maps from the NeuroSynth database (*** *P* < 0.001, FDR-corrected). For **b**, serotonin transporter (5-HTT), receptor (5-HT_1A_, 5-HT_1B_, 5-HT_2A_, 5-HT_4_, 5-HT_6_). Dopamine transporter (DAT), receptor (D_1_, D_2_). Norepinephrine transporter (NET). Histamine receptor (H_3_). Acetylcholine transporter (VAChT), receptor (M_1_, α_4_β_2_). Cannabinoid receptor (CB_1_). Opioid receptor (MOR). Glutamate receptor (mGluR_5_). GABA receptor (GABA_A/BZ_).

## The glioma network is associated with action

To characterize the functional properties of the glioma network, we matched the network to 1,334 functional activation patterns within the Neurosynth meta-analysis database^53^. After excluding purely anatomical functional patterns, we generated a word cloud highlighting behavior-related functional patterns that were significantly correlated with the glioma network (Pearson’s *r* > 0.100, *P* < 0.001, FDR-corrected; Supplementary Fig. 4). Functional maps of the eight most significantly correlated patterns all involved action-related neural systems (Fig. 5c), which can be organized into three types. The first type centered on action-modulatory feedback processing, exemplified by ‘pain’ (Pearson’s *r* = 0.430, CI = [0.408, 0.451]) and ‘noxious’ (Pearson’s *r* = 0.276, CI = [0.252, 0.300]). The second type comprised core motor execution components: ‘motor’ (Pearson’s *r* = 0.319, CI = [0.295, 0.343]), ‘speech production’ (Pearson’s *r* = 0.282, CI = [0.258, 0.306]) and ‘movement’ (Pearson’s *r* = 0.272, CI = [0.247, 0.296]). The final type included sensory-motor integration systems, such as ‘somatosensory’ (Pearson’s *r* = 0.365, CI = [0.342, 0.388]), ‘sensorimotor’ (Pearson’s *r* = 0.299, CI = [0.275, 0.323]) and ‘auditory’ (Pearson’s *r* = 0.284, CI = [0.260, 0.308]) processing domains. These results demonstrate that the glioma network engages a functional circuitry essential for action initiation, execution and feedback.

## Discussion

### AMN as a highly active and functional substrate for glioma growth

The emerging field of cancer neuroscience represents a paradigm shift, moving from an exclusive focus on tumors to an exploration of the intricate interactions between gliomas and the brain^60–68^. Recent studies demonstrated that extensive neurotransmitter-mediated glioma-neuron crosstalk establishes permissive neural circuits that promote tumorigenesis and progression^21–27,69^. Glioma cells could rapidly integrate into long-range functional circuits even in contralateral cortex^21,70^. It is not surprising that it can induce widespread functional alterations not only in the tissue adjacent to the lesion but also in distant brain regions^71,72^. The body of evidence supports the notion that glioma should be regarded as a neurological disease affecting the entire brain, and gliomas, in turn, may be affected by functional activities^18,65,73,74^.

By leveraging RSFC, we identified the AMN as a critical functional nexus for glioma pathogenesis. This association may be driven by three interrelated mechanisms. First, as a hub of action-related neural processing, the AMN exhibits tonically elevated metabolic demand, while awake^32,33^. Active neurons in this network release activity-dependent neurotransmitters such as acetylcholine, dopamine, and serotonin^21–24,27^. Glioma cells express receptors for these neurotransmitters, which can promote a more motile phenotype conducive to glioma growth^21,23,27,75^. Second, the brain exhibits significant inter-regional heterogeneity in cellular composition and associated immune landscapes^76–78^. Notably, neuronal activity dynamically regulates microglial function via norepinephrine signaling^79^. In awake state, microglia display reduced arborization and restricted surveillance territories compared to anesthetized conditions. Elevated neuronal firing suppresses microglial motility and immune surveillance^80^. This suggests that the AMN, as a hub of sustained activity during wakefulness, may exhibit heightened immunosuppressive microenvironments. Third, task-evoked neural activity in the AMN may induce transcriptional changes. Neuronal activity-dependent secretion of neuroligin-3 (NLGN3) has been shown to upregulate oncogenic pathways, including the proto-oncogene FOS, and to enhance NLGN3 expression within glioma cells^11,12,81,82^. Concurrently, acetylcholine accumulation in the AMN can trigger the expression of immediate early genes (e.g., FOS and FOSB, EGR1), nuclear transcription factors associated with Ca^2+^ oscillations and epigenetic regulators^21^. These mechanisms suggest that the AMN activity may contribute to the aberrant transcription of oncogenes, potentially acting in concert with pre-existing driver mutations to promote glioma initiation and progression.

### AMN dysfunction as a mechanism underlying glioma-related symptoms

We identified that the glioma network mainly supported action-related processing (Fig. 5c), from sensory-motor integration, movement execution to behavioral adaptation through feedback. This functional property provides a neural substrate explaining the prevalence of action-related clinical symptoms in glioma patients. Notably, our findings demonstrate striking convergence with the 2020 US National Cancer Institute proposed core patient-reported outcomes for gliomas^83^; these clinical markers, e.g., communication deficits, aphasia, and physical dysfunction, show alignment with functional domains identified in our analyses including ‘auditory’, ‘speech production’, and ‘motor’ (Fig. 5a). In addition, approximately 25% of glioma patients report chronic bodily pain^84^, potentially stemming from glioma-induced dysfunction of pain-processing pathways. Beyond pure symptoms (e.g., aphasia and motor deficits), our results shed light on the frequent neuropsychiatric comorbidities (weakness, fatigue, apathy and abulia)^85,86^ that conventional lesion models struggle to explain. We propose these manifestations may represent network-level dysfunction of the AMN, the principal neural substrate for sustaining high-arousal states essential for action^32^. Intriguingly, our model also offers a novel pathophysiological mechanism for the poorly understood prevalence of axial somatic symptoms (nausea, dyspnea, constipation) in glioma populations^84^. The AMN-connected SCAN^41^, governing axial movement coordination, may undergo glioma-mediated disorganization, potentially disrupting the ability of gastrointestinal motility, respiratory modulation, and visceral sensory processing.

### AMN as a therapeutic intervention target

Glioma cells exhibit functional integration into neural circuits by receiving and processing diverse neurotransmitter signals^21–27^. While glutamatergic synaptic connectivity between neurons and glioma cells has been well-documented^14^, recent studies have identified cholinergic synapses as a critical neuron-glioma communication interface^21–24,59^. Specifically, ACh signaling through muscarinic receptors (e.g., CHRM3) promotes tumor progression by enhancing migratory capacity and inducing transcriptional reprogramming. Our spatial mapping strengthens this mechanistic link, demonstrating a significant positive correlation between the distribution of VAChT and the glioma network topology (Fig. 5a). Importantly, preclinical studies demonstrate that CHRM3 knockdown suppresses glioma migration and extends survival in murine models^21–24^, positioning cholinergic signaling as a promising therapeutic target. In addition, we observed spatial enrichment of dopamine and serotonin transporter clusters within the AMN (Fig. 5b). It showed concordance with previous studies that report a reduced glioblastoma incidence and improved survival outcomes in schizophrenia patients receiving antipsychotic therapy targeting dopamine and serotonin receptors^87,88^. These findings suggest that, beyond cholinergic synapses, dopamine and serotonin signaling in the AMN may serve as actionable therapeutic targets in glioma treatment, warranting further systematic investigation.

Recently, emerging nonsurgical and nonpharmacologic interventions, such as tumor-treating fields^89–91^ and chimeric antigen receptor T-cell therapy^92–97^, have been evaluated as adjuncts to standard glioma treatments (surgery, radiation and chemotherapy). Most of these approaches primarily focus on direct tumor targeting^90,92^. Here, the association between the AMN and glioma distribution presents a new therapeutic premise: modulation of neuronal activity within the AMN—even within regions distant from the tumor—may inhibit glioma progression by suppressing neuron-glioma synaptic communication. This network-based therapeutic rationale suggests that noninvasive brain stimulation techniques could offer therapeutic potential by altering synaptic transmission. A preliminary clinical investigation employing transcranial direct current stimulation in glioma patients demonstrated significant hemodynamic alterations, evidenced by a 36% reduction in intratumoral cerebral blood flow (CBF)^98^. While this intervention targeted tumor tissue directly, it provided crucial preliminary evidence regarding the feasibility and safety of neuromodulation in glioma. Given the established efficacy of neuromodulation techniques in altering functional connectivity^99–106^, strategic targeting of AMN, rather than focusing solely on tumor mass, may suppress neuron-glioma interactions and ultimately slow tumor growth.

---

**Extended Data Fig. 1.**
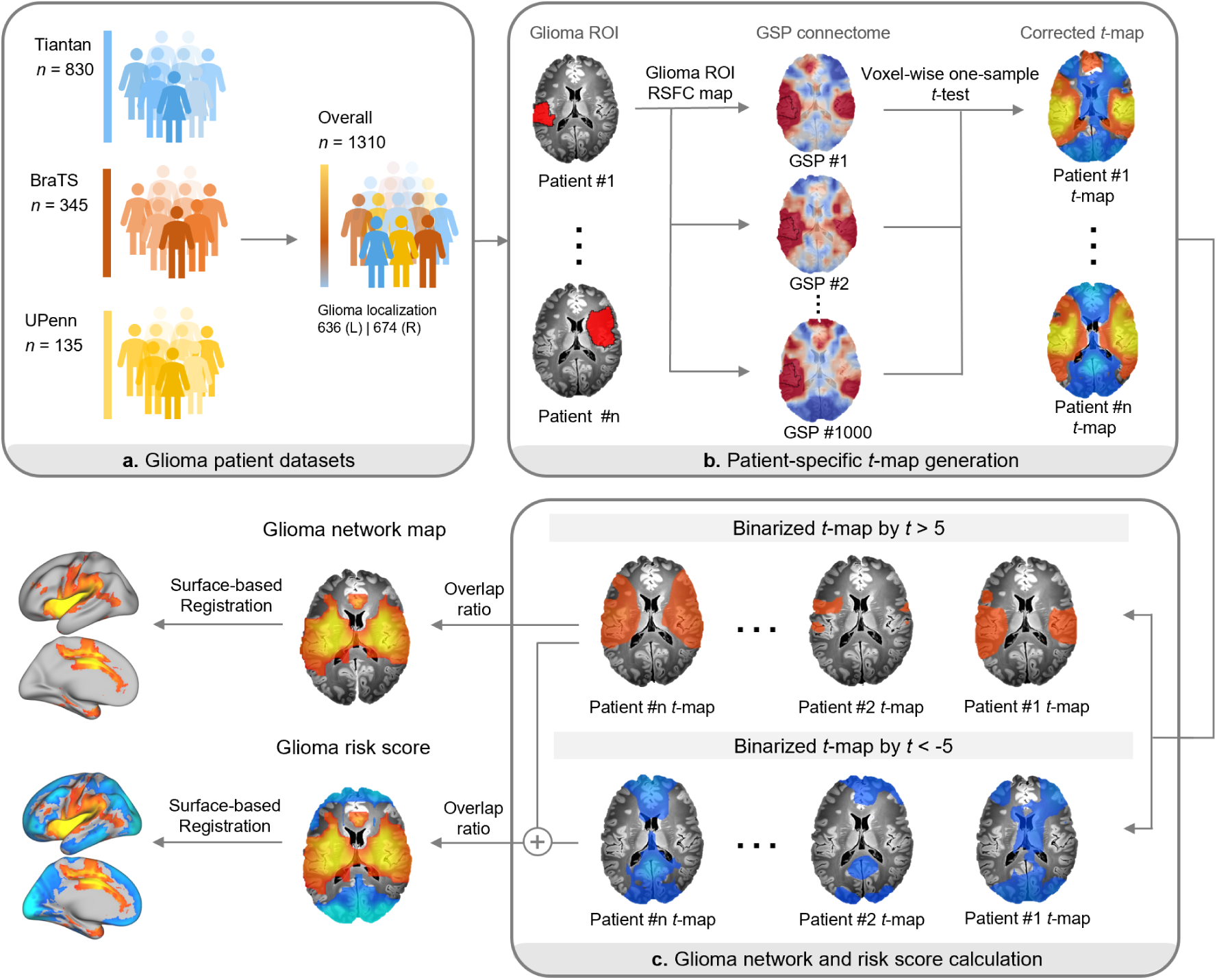
Framework of the glioma network mapping algorithm. **a,** Glioma network mapping was performed using three independent datasets: Tiantan (*n* = 830), BraTS (*n* = 345)^107–110^, and UPenn (*n* = 135)^111^, which were combined into an overall dataset (*n* = 1,310). **b,** For each patient’s glioma region of interest (ROI), resting-state functional connectivity (RSFC) was computed using rs-fMRI data from 1,000 healthy participants in the Brain Genomics Superstruct Project (GSP) connectome^112^. The average BOLD (blood-oxygenation level dependent) signal time series within the glioma ROI was correlated with the time series of all other brain voxels, and the resulting correlation coefficients were transformed to *z*-scores using Fisher’s *r*-to-*z* transformation. Voxel-wise one-sample *t*-tests were then performed across all RSFC maps, and FDR correction (*P* < 0.01) was applied to generate patient-specific *t*-maps. **c,** To define the glioma network map, each patient-specific *t*-map was thresholded at |*t*| > 5 and binarized to identify regions of significant positive (*t* > 5) and negative (*t* < -5) connectivity. The overlap ratio of positive connectivity across patients was computed to construct the glioma network. Negative connectivity was also included and combined with positive results to calculate the glioma risk score, which quantifies the likelihood that a given voxel is anti-correlated with glioma-prone regions. A higher positive score indicates a greater likelihood of glioma occurrence in that voxel. The final maps were registered to the fsaverage6 cortical surface for visualization.

**Extended Data Fig. 2.**
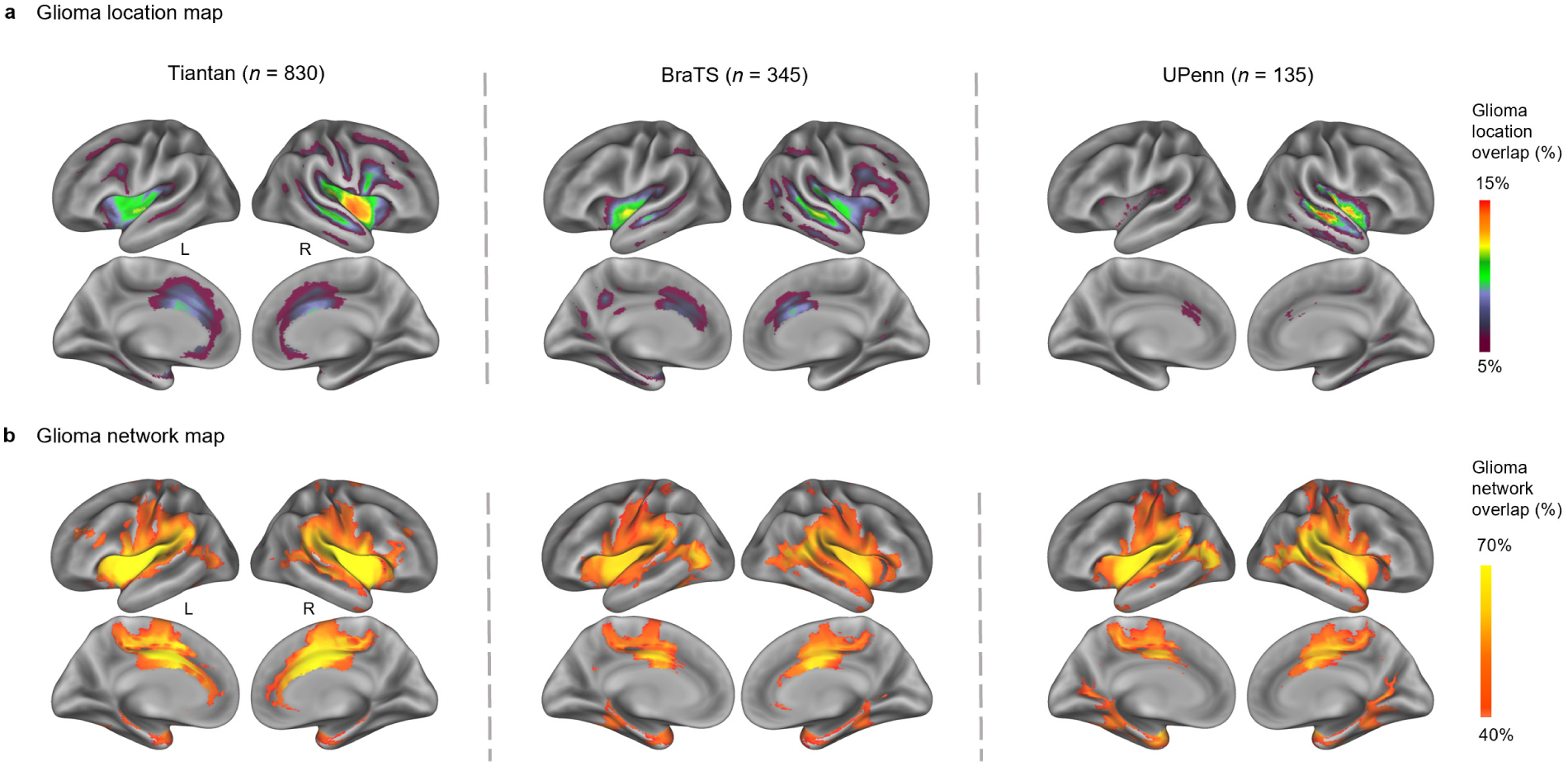
Surface maps of glioma location and network. **a,** Whole-brain surface maps of glioma anatomical locations from three independent datasets: Tiantan (*n* = 830), BraTS (*n* = 345), and UPenn (*n* = 135). The color in the glioma location map represents the cross-individual anatomical overlap of the gliomas. **b,** Whole-brain surface maps of glioma networks derived from glioma network mapping for each dataset (see Methods). The color in the glioma network map represents the cross-individual functional overlap of the gliomas.

**Extended Data Fig. 3.**
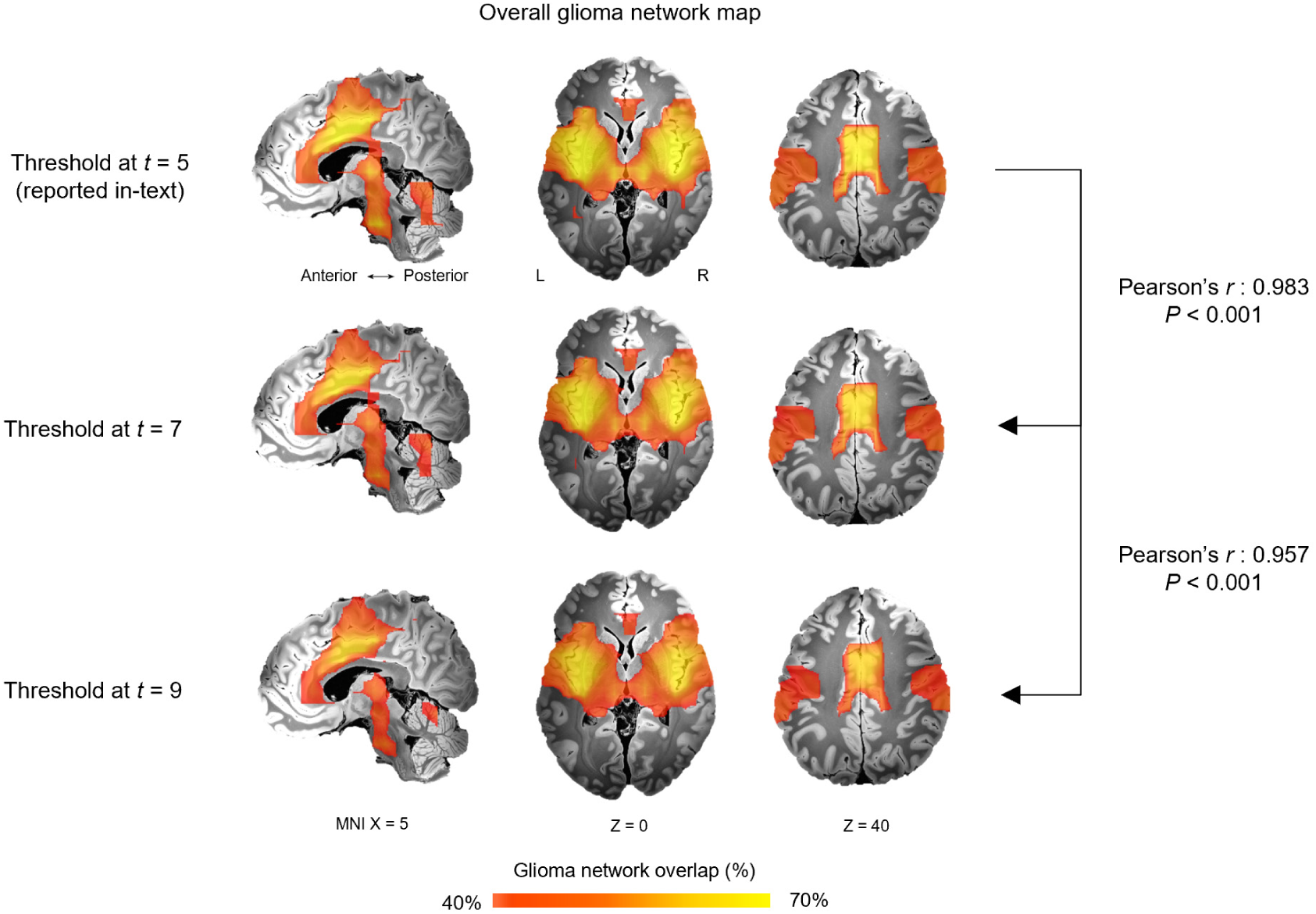
Overall glioma network maps across different threshold cut-offs. Patient-specific *t*-maps were binarized using three different positive connectivity thresholds (*t* = 5 [reported in-text], *t* = 7, and *t* = 9). For each threshold, functional overlaps were computed across patients within our glioma dataset to generate the overall glioma network map (*n* = 1,310; see Methods). The resulting glioma network maps showed high spatial consistency across thresholds (Pearson’s *r* = 0.983 between *t* = 5 and *t* = 7, *P* < 0.001, CI = [0.982, 0.984], uncorrected; Pearson’s *r* = 0.957 between *t* = 5 and *t* = 9, *P* < 0.001, CI = [0.955, 0.959], uncorrected).

**Extended Data Fig. 4.**
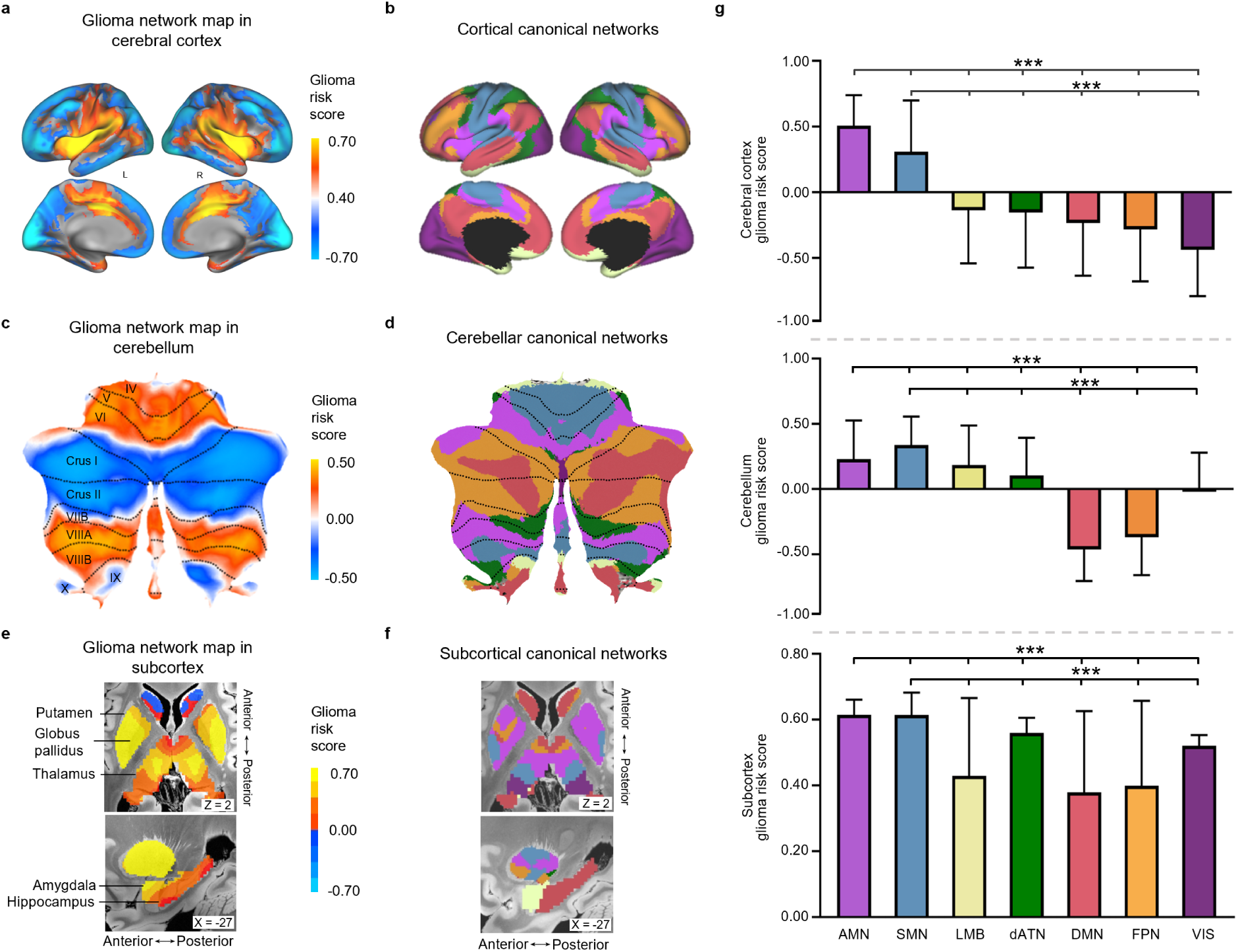
Distribution of glioma risk scores across canonical networks in cortex, cerebellum and subcortex. **a,** Glioma risk scores mapped to the cerebral cortex, defined as the cross-participant overlap of glioma network connectivity (positive values) and anti-correlations (negative values). **b,** resting-state functional connectivity (RSFC)-derived seven-network atlas of the cerebral cortex^54^. **c,** Glioma risk scores in the cerebellum (flat map). **d,** RSFC-derived seven-network atlas of the cerebellum^55^. **e,** Glioma risk scores in subcortical regions. **f,** RSFC-derived seven-network atlas of the subcortex^56,57^. **g,** Mean glioma risk scores across canonical functional networks in the cortex (top), cerebellum (middle), and subcortex (bottom). In the cortex, the action-mode network (AMN) shows the highest glioma risk score, followed by the somatomotor network (SMN). In the cerebellum, the SMN shows the highest score, followed by the AMN. In the subcortex, the AMN exhibits the highest score, followed by the SMN. All comparisons are based on two-tailed independent *t*-tests versus AMN or SMN (\*\*\**P* < 0.001, FDR-corrected). Full values for each network are provided in Supplementary Table 5. For **g**, action-mode network (AMN). Somatomotor network (SMN). Limbic Network (LMB). Dorsal attention Network (dATN). Default-mode Network (DMN). Frontoparietal network (FPN). Visual Network (VIS).

**Extended Data Fig. 5.**
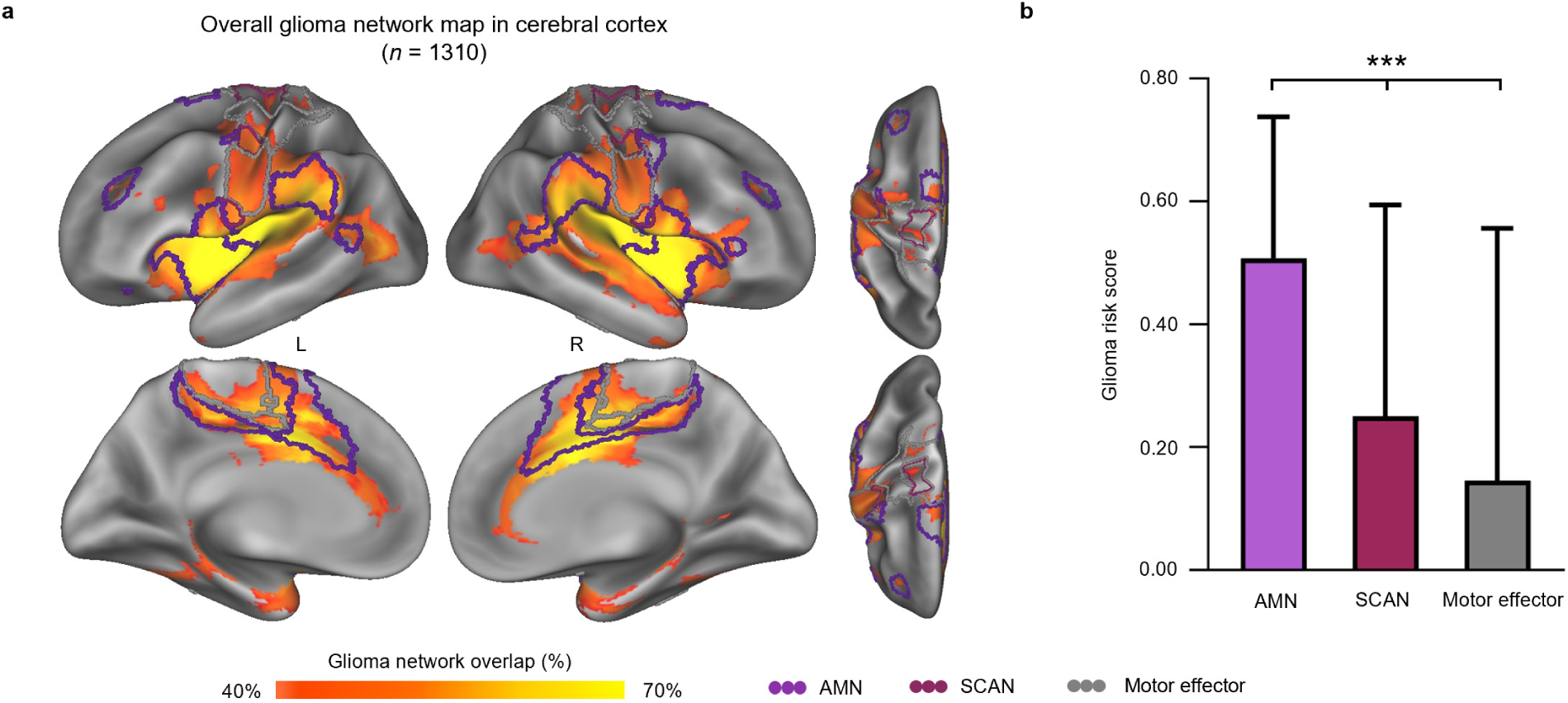
Distribution of glioma network map across action-mode network (AMN), somato-cognitive action network (SCAN), and motor effector network. **a,** Surface-based glioma network map overlaid on the cerebral cortex. The AMN is outlined in purple, the SCAN in red, and the motor effector network (including foot, hand, and mouth regions) in grey. **b,** Glioma risk scores averaged across these cortical networks. The AMN (0.508 ± 0.230) exhibits the highest glioma risk score, followed by the SCAN (SCAN: 0.251 ± 0.344; motor effector network: 0.146 ± 0.414; two-tailed independent t-tests with AMN or SCAN, *** *P* < 0.001, FDR-corrected).

**Extended Data Fig. 6.**
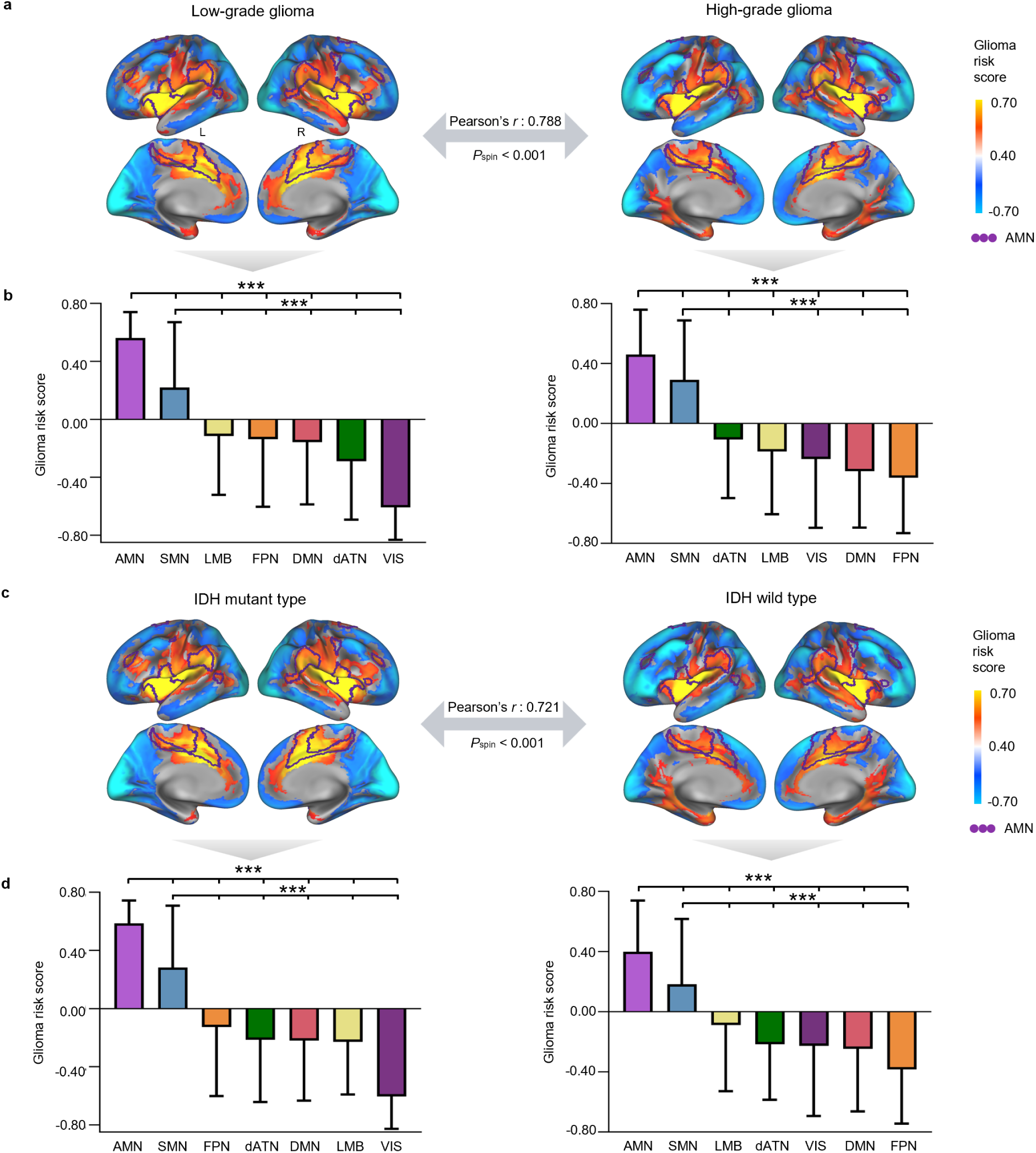
Similarity of glioma network maps across glioma grades and IDH (isocitrate dehydrogenase) subtypes. **a,** Spatial similarity between low- and high-grade glioma network maps derived from the Tiantan dataset (*n* = 830), with statistical significance evaluated using the spin test. **b,** Box plots showing the glioma risk score of glioma network maps within the cortical functional networks for low-grade (*n* = 432, age = 38.9 ± 12.2 years) and high-grade groups (*n* = 398, age = 49.5 ± 15.2 years). Among all networks, the AMN (0.572 ± 0.178, low-grade; 0.470 ± 0.298, high-grade) exhibits the highest functional overlap, followed by the SMN (low-grade: 0.222 ± 0.450; high-grade: 0.294 ± 0.395). **c,** Spatial similarity between IDH mutant and wild type glioma network maps derived from the Tiantan dataset (*n* = 830), with statistical significance assessed using the spin test. **d,** Box plots showing the glioma risk score of glioma network maps within the cortical functional networks for IDH-mutant (*n* = 477, age = 41.7 ± 12.6 years) and IDH wild (*n* = 353, age = 47.0 ± 16.7 years). Similar to grade subgroups, the AMN (mutant type: 0.599 ± 0.155; wild type: 0.403 ± 0.340) demonstrates the highest functional overlap, followed by the SMN (mutant type: 0.285 ± 0.423; wild type: 0.187 ± 0.430). Statistical comparisons for AMN or SMN were performed using two-tailed independent t-tests (all *** *P* < 0.001, FDR-corrected). The detailed values of the glioma risk scores in the seven networks of cerebral cortex, cerebellum and subcortex are given in Supplementary Table 5.

**Extended Data Fig. 7.**
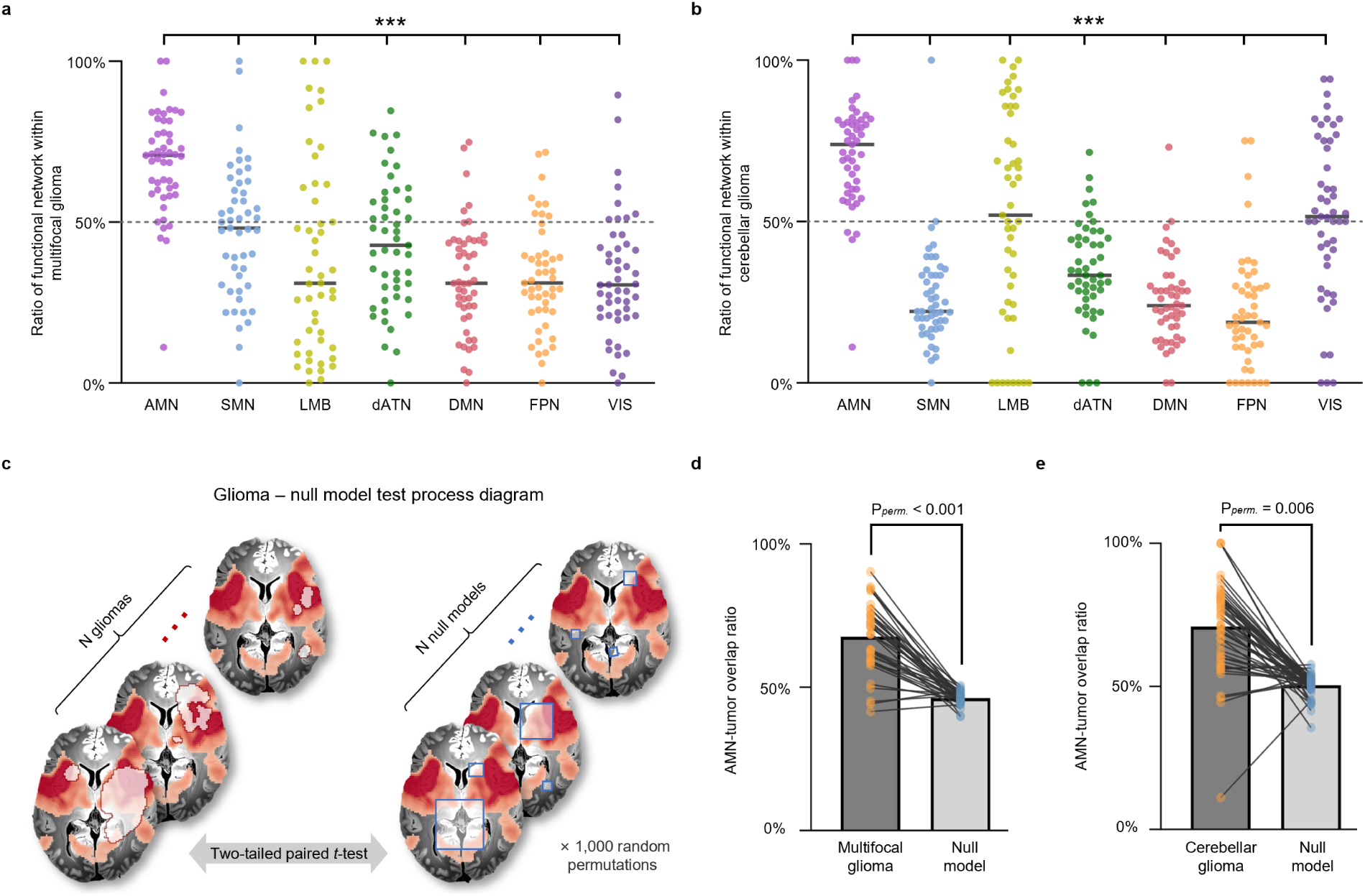
Network comparison and permutation tests for multifocal and cerebellar gliomas. **a,** Ratios of canonical functional network regions located within multifocal gliomas. Each dot represents one patient; dashed line marks 50% network coverage within tumor regions. Across networks, tumors exhibited significantly greater overlap with the action-mode network (AMN) than with any other canonical network (two-tailed paired *t*-test, all *P* < 0.001, FDR-corrected). **b,** Ratios of canonical functional network regions located within cerebellar gliomas. Tumor locations showed significantly stronger alignment with the AMN than with any other network (two-tailed paired *t*-test, all *P* < 0.001, FDR-corrected). **c,** Process of the glioma-null model permutation test. Each of the *N* gliomas was paired with *N* randomly placed null models with identical size and number. A two-tailed paired *t*-test was conducted between gliomas and their corresponding null models, iterating 1,000 times to assess statistical significance. **d,** The AMN-tumor overlap ratios of 38 multifocal glioma patients (orange) were compared to the mean AMN-tumor overlap ratios of their 38 null models (blue) across 1,000 permutations (multifocal gliomas: 68.9% ± 13.2% versus null model: 47.4% ± 1.9%, *P*_perm._ < 0.001). **e,** The AMN-tumor overlap ratios of 49 cerebellar glioma patients (orange) were compared to the mean AMN-tumor overlap ratios of their 49 null models (blue) across 1,000 permutations (cerebellar glioma: 71.0% ± 6.0% versus null model: 50.4% ± 4.0%; *P*_perm._ = 0.006). Each line represents one patient.

## Methods

### Patient demographics and clinical characteristics

This study included five independent datasets comprising a total of 1,397 glioma patients. The demographic and clinical characteristics of these patient samples are presented in Supplementary Table 1.

The Tiantan, Cerebellar glioma and Multifocal glioma datasets were retrospectively collected from patients at the Tiantan Hospital, Capital Medical University, China, between 2015 and 2023, with no overlapping patients across the datasets. Patients in these three cohorts underwent routine preoperative MRI, including T1-weighted imaging (T1w), contrast-enhanced T1-weighted contrast-enhanced imaging (T1-CE), T2-weighted imaging (T2w), and T2-weighted fluid-attenuated inversion recovery imaging (T2-FLAIR). The Tiantan dataset included patients with single cortical gliomas, while the Cerebellar and Multifocal Glioma datasets comprised those with gliomas in the cerebellum and multiple brain regions, respectively. Inclusion criteria were as follows: 1) histological confirmation of gliomas according to the 2021 World Health Organization classification criteria^113^; 2) availability of routine preoperative MRI data. Exclusion criteria included: 1) history of prior brain tumor surgery; 2) inability to undergo MRI scanning. Informed consent was obtained from all participants, and the retrospective analysis was approved by the Institutional Review Board of Tiantan Hospital.

In addition, two publicly available datasets were included in this study, for which preprocessed glioma segmentation masks were provided. The BraTS dataset, obtained from the Multimodal Brain Tumor Image Segmentation Challenge 2021 (http://braintumorsegmentation.org)^107–110^, consisted of 345 cortical glioma patients from multiple institutions. The UPenn dataset, sourced from the University of Pennsylvania Health System (https://brain.labsolver.org/upenn_gbm.html)^111^, included 145 patients diagnosed with cortical glioblastoma. Due to failed image registration in 10 cases, a total of 135 patients were included in our analysis.

### Glioma location overlap

Gliomas in the Tiantan, Cerebellum and Multifocal datasets were manually segmented on T2-weighted fluid-attenuated inversion recovery (T2-FLAIR) MRI scans by two board-certified neuroradiologists using MRIcroN v1.0^114^. Segmentation was based on consensus readings, following protocols established in our previous study^20^. Regions with abnormal hyperintense signals on T2-FLAIR images were identified as glioma tissue. These initial segmentations were subsequently reviewed and refined by a senior neuroradiologist to define definitive glioma boundaries. For the BraTS and UPenn datasets, glioma masks were segmented based on multi-modal imaging, including T1w, T1-CE, T2w, and T2-FLAIR scans^107–111^. BraTS masks were publicly available, generated by fusing top-performing algorithms from previous BraTS challenges ^115–118^, and manually refined by board-certified neuroradiologists. For the UPenn dataset, glioma masks were segmented automatically using BraTS challenge algorithms^115–118^ and visually inspected, with manual corrections applied as required.

To investigate the spatial distribution of gliomas across specific brain regions, all glioma masks were mapped to a 1-mm spatial resolution volumetric template (the FSL-version of the MNI ICBM152 nonlinear template) with a 12-degree-of-freedom linear transformation (FSL v6.0)^119^. The standardized glioma masks were then merged to generate a spatial overlap heatmap of gliomas for each dataset, where each voxel represents the frequency of glioma occurrence. These voxel-based maps were finally registered to the surface-based fsaverage standard space using FreeSurfer.

### Glioma network mapping

We adapted lesion network mapping^46–50^ and proposed a glioma network mapping approach (Extended Data Fig. 1) to investigate the functional correlations between gliomas with specific brain networks. This approach enables quantification of the brain-wide functional connectivity of glioma locations using normative resting-state functional MRI (rs-fMRI) data from 1,000 healthy individuals from the Brain Genomics Superstruct Project (GSP)^112^. Rs-fMRI data were preprocessed using DeepPrep^120–122^, with the same parameter settings as in our previous studies^123,124^. Each glioma lesion was defined as a region of interest (ROI). For each healthy participant, the mean BOLD time series within the glioma ROI was extracted and correlated with the time series of every other brain voxel, yielding individual RSFC maps. These correlation coefficients were then normalized using Fisher’s *r*-to-*z* transformation. Voxel-based one-sample *t*-tests were then performed on the *z*-values across all 1,000 participants, and FDR correction (*P* < 0.01) was applied to the *z*-values to generate a glioma ROI-specific *t*-map. The *t*-values quantified the strength and consistency of the functional connectivity between the glioma and the rest of the brain.

We then thresholded the corrected maps at |*t*| > 5 to binarize the functional connections: voxels with *t* > 5 were defined as positively connected to the glioma, while those with *t* < -5 were defined as negatively connected (anti-correlated). We then computed the overlap ratio of positive functional connectivity across all patients to derive the glioma network. To better quantify the network’s functional subdivisions, we also calculated the overlap ratio of negative functional connectivity. Both the positive connectivity and anti-correlations of the glioma network were integrated to generate the glioma risk score. A higher positive score indicates a greater likelihood of glioma occurrence in that voxel, while a higher negative score suggests a lower probability. To assess the robustness of our findings, we repeated the analysis using higher threshold values (|*t*| > 7 and |*t*| > 9) and evaluated result consistency via spatial Pearson correlation coefficients.

### Correlation with distinct brain networks

To investigate whether the glioma network preferentially aligns with specific functional brain networks, we mapped the constructed glioma network onto a canonical resting-state functional connectivity (RSFC)-derived atlas encompassing the cerebral cortex, cerebellum, and subcortex. The cortical, cerebellar, and subcortical parcellations were defined using publicly available atlases: Yeo et al. (2011)^54^ for the cortex, Buckner et al. (2011)^55^ for the cerebellum, Choi et al. (2012)^56^ for the striatum atlas. We then analyzed the distribution of glioma risk scores within each of the seven networks separately for the cortex, cerebellum, and subcortex, thereby obtaining the whole-brain distribution of glioma risk scores.

These functional networks encompass the visual network (VIS), somatomotor network (SMN), dorsal attention network (dATN), AMN, frontoparietal network (FPN), default-mode network (DMN), and limbic network (LMB). The AMN has been proposed as a reconceptualization of the ventral attention network (vATN), emphasizing its role in goal-directed behavior^32^. For the subcortex, the original seven-network atlas provided coverage for only the striatum. To ensure a more comprehensive characterization, based on a topographic atlas derived from functional gradients^57^, we used rs-fMRI data from 1,000 healthy participants in the GSP^112^, and assigned the thalamus, hippocampus, amygdala and globus pallidus to the most functionally relevant cortical network through a winner-take-all classification based on its highest correlation with the mean signal of the cortical seven networks. To mitigate potential signal contamination from adjacent cortical areas, subcortical signals were regressed out to remove contributions from immediately adjacent cortical regions. For cerebellar glioma network maps, we registered them onto SUIT space and visualized them using the SUITPy toolbox^125^.

### Predicting the occurrence of multifocal and cerebellar gliomas

To assess the preferential localization of gliomas within the AMN, we applied this network to predict glioma occurrence in independent datasets of multifocal gliomas (*n* = 38) and cerebellar gliomas (*n* = 49). Normative resting-state functional connectivity (RSFC) pattern of the AMN was used as a spatial predictor. For each patient, the overlap ratio between the tumor mask and the AMN was calculated as the proportion of tumor voxels residing within the network relative to the total tumor volume. We also compared the prediction with those derived from six other canonical functional networks (sensorimotor [SMN], limbic [LMB], dorsal attention [dATN], default mode [DMN], frontoparietal [FPN], and visual [VIS]). The resulting overlap ratios across networks were compared using two-tailed paired *t*-tests, and *P*-values were corrected for multiple comparisons using the false discovery rate (FDR) method. To further determine whether glioma localization within the AMN exceeded chance levels, we employed a glioma-null model permutation test with 1,000 random permutations on the multifocal and cerebellar glioma datasets, respectively. Specifically, we compared the AMN-tumor overlap ratio for actual glioma masks against those of randomly positioned, null model-based ROIs within the cerebral or cerebellar regions. To ensure the robustness of the results, the null model-based ROIs were volume-matched to the corresponding glioma masks, and the number of tumors was matched identically to those observed in multifocal gliomas. The statistical significance of the permutation test was assessed by performing a two-tailed paired *t*-test for each iteration of the glioma-null model. The overall significance was determined by the proportion of iterations yielding a *P*-value < 0.05.

### Predicting survival times in glioma patients

We performed survival analysis using two independent datasets: 164 patients from the Tiantan dataset and 121 patients from the UPenn dataset, both of which contained explicit overall survival data and censoring information (Supplementary Table 6). From the original UPenn dataset (*n* = 135), two patients were excluded due to missing survival or censoring information. An additional 12 patients aged over 80 years were excluded, considering the established prognostic role of age in glioblastoma^126^ and general life expectancy in the U.S.^127^. This yielded a final sample of 121 patients for analysis.

To stratify patients into two groups, we calculated the AMN-FC-weighted overlap score for each patient. This score quantifies the extent of AMN-tumor overlap, weighted by the voxel-wise strength of AMN RSFC within the overlapping region. Using maximally selected rank statistics^58^, we determined the optimal stratification threshold (cutoff = 0.10) to dichotomize patients into high-overlap and low-overlap groups. This non-parametric approach systematically identifies the threshold that maximizes the separation of survival distributions based on log-rank statistics. Kaplan-Meier survival analyses were then performed separately for each dataset, and group differences were evaluated using two-sided log-rank tests. To assess the prognostic relevance of the overlap score relative to established clinical and molecular factors (age, sex, tumor volume, glioma grade, IDH mutation status, and MGMT promoter methylation status), we conducted both univariate and multivariate Cox proportional hazards regression analyses.

### Spin-based null modeling of glioma-neurotransmitter spatial correlations

We leveraged the Neuromap toolbox (https://netneurolab.github.io/neuromaps/) to access density maps for 18 neurotransmitter receptors and transporters derived from PET tracer studies^51,52^. These maps correspond to nine key neurotransmitter systems, including dopamine (D_1_, D_2_, DAT), noradrenaline (NET), serotonin (5-HT_1A_, 5-HT_1B_, 5-HT_2A_, 5-HT_4_, 5-HT_6_, 5-HTT), acetylcholine (α_4_β_2_, M_1_, VAChT), glutamate (mGluR_5_), GABA (GABA_A/BZ_), histamine (H_3_), cannabinoid (CB_1_), and opioid (MOR). All density maps were registered to a cerebral atlas comprising 213 fine-grained functional regions (see Supplementary Fig. 5 for the detailed atlas)^112,128,129^. Spatial similarity between the glioma network map and each neurotransmitter receptor/transporter distribution was then calculated using the Pearson correlation coefficient. To control for spatial autocorrelation, we performed rotation-based null model (spin) permutation testing via the BrainSpace toolbox^130^. Vertex-wise glioma network maps were aligned to the fsaverage6 surface template and rotated 10,000 times per hemisphere, with vertices rotated into the medial wall excluded. These permutations preserved cortical spatial autocorrelation and topology while randomizing anatomical alignment. Following each rotation, the spun glioma network map was re-parcellated into a 213-region cortical atlas^112,128,129^ to calculate correlations with each neurotransmitter density map. Statistical significance was defined as the proportion of permuted correlations (|*r*_spin_|) that exceeded the observed correlation (|*r*_observation_|). *P*-values were corrected for multiple comparisons across all 18 neurotransmitter systems using the false discovery rate (FDR) method. Tracer references and full statistical results are provided in Supplementary Table 8.

### Comparative functional pattern analytics

To characterize the cognitive and behavioral correlates of the glioma network, we conducted a spatial meta-analytic comparison using the NeuroSynth database (https://neurosynth.org/)^53^. Neurosynth provides whole-brain activation maps for 1,334 functional patterns, which are derived from large-scale meta-analyses of neuroimaging studies. These maps represent how fluctuations in regional brain activity correspond to specific psychological processes. In our analysis, we excluded anatomical (e.g., ‘insula’ and ‘cortical’) and repetitive patterns, focusing exclusively on cognitive and behavioral correlates. Voxel-wise Pearson correlation coefficients were used to quantify the spatial similarity between the glioma network map and each functional spatial phenotype map.

## Data availability

The BraTS and UPenn datasets used in this study are publicly available at http://braintumorsegmentation.org and https://brain.labsolver.org/upenn_gbm.html, respectively. All patient data supporting the findings of this study are available from the corresponding websites upon reasonable request. The Tiantan, multifocal, and cerebellar glioma datasets are available under restricted access, in accordance with the policies and procedures of Tiantan Hospital, Beijing, China. Data access requests can be made by contacting Y.W. The normative rs-fMRI data from GSP used for functional connectivity analyses are publicly available at https://dataverse.harvard.edu/dataverse/GSP. The cortical, cerebellar, and striatum parcellation atlases used for network-level analyses are publicly available online, including the Yeo et al. (2011) cortical parcellation (https://surfer.nmr.mgh.harvard.edu/fswiki/CorticalParcellation_Yeo2011), the Buckner et al. (2011) cerebellar parcellation (https://surfer.nmr.mgh.harvard.edu/fswiki/CerebellumParcellation_Buckner2011), and the Choi et al. (2012) striatum parcellation (https://surfer.nmr.mgh.harvard.edu/fswiki/StriatumParcellation_Choi2012). The volumetric brain template is an ultrahigh-resolution ex-vivo brain in MNI space^131^, which is available at https://datadryad.org/stash/downloads/file_stream/182489.

## Code availability

The codes for the glioma network mapping algorithm and other analyses in this study are available on GitHub at https://github.com/pBFSLab/Glioma_AMN. Software packages incorporated into the above code for data analysis included: Matlab R2020a, https://www.mathworks.com/; Python v3.11.6, https://www.python.org; MRIcroN v1.0, https://www.nitrc.org/projects/mricron; Connectome Workbench v1.5, http://www.humanconnectome.org/software/connectome-workbench.html; Freesurfer v6.0.0, https://surfer.nmr. mgh.harvard.edu/; FSL v6.0, https://fsl.fmrib.ox.ac.uk/fsl/fslwiki; SUITPy toolbox, https://github.com/DiedrichsenLab/SUITPy.

## Acknowledgment

This work was supported by the Changping Laboratory (H.L.); National Institutes of Health grants MH096773 (N.U.F.D.), MH122066 (E.M.G., N.U.F.D.), MH121276 (E.M.G., N.U.F.D.), MH124567 (E.M.G., N.U.F.D.), NS129521 (E.M.G., N.U.F.D.), and NS088590 (N.U.F.D.); the Beijing Natural Science Foundation of China JQ23040 (Y.W), L241027 (Y.W.); the Intellectual and Developmental Disabilities Research Center (N.U.F.D.); by the Kiwanis Foundation (N.U.F.D.); and the Washington University Hope Center for Neurological Disorders (E.M.G., N.U.F.D.).

## Author contributions

Conception: H.L., J.R. and W.C. Design: W.C., J.Z., Z.Y. and J.R. Data acquisition, analysis and interpretation: W.C., J.Z, Z.Y., T.J., H.B., S.F., Z.C., X.F., E.M.G., Y.W. and H.L. Manuscript writing and revision: W.C., J.Z., E.M.G., S.M., V.M.S., S.S., D.W., N.U.F.D., and H.L. All authors approved the final manuscript.

