## Supplementary Materials for "Gliomas preferentially develop within the action-mode network"

### Online Supplementary Materials

**Supplementary Fig. 1 | Glioma functional connectivity and its anti-correlations.**
**Supplementary Fig. 2 | Full-threshold display of the cortical and subcortical distribution of the glioma network.**
**Supplementary Fig. 3 | Cutoff optimization for AMN-tumor overlap score.**
**Supplementary Fig. 4 | Comparative functional pattern analytics based on the NeuroSynth database.**
**Supplementary Fig. 5 | Parcellation of 213 brain functional regions.**
**Supplementary Table 1 | Patient demographic and clinical data.**
**Supplementary Table 2 | Specific coordinates of glioma location foci across datasets.**
**Supplementary Table 3 | Specific coordinates of glioma network foci across datasets.**
**Supplementary Table 4 | Specific coordinates of overall glioma network foci.**
**Supplementary Table 5 | Glioma risk scores across canonical networks.**
**Supplementary Table 6 | Demographics and clinical summary of patient datasets used for survival analysis.**
**Supplementary Table 7 | Univariate and multivariate survival analysis of patient datasets.**
**Supplementary Table 8 | Details of neurotransmitters and other metabolism measures datasets.**

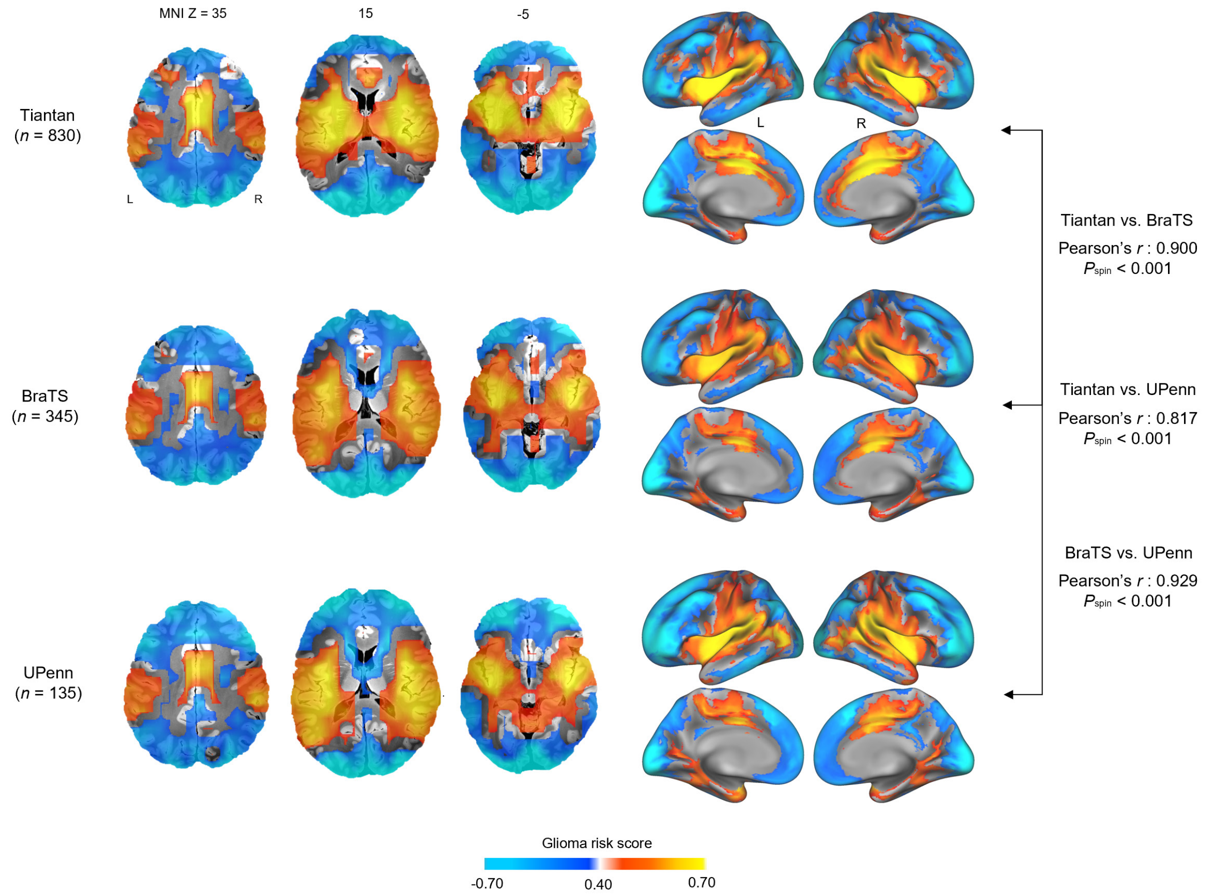


**Supplementary Fig. 1 | Glioma functional connectivity and its anti-correlations.** Glioma network maps and their corresponding anti-correlations, derived from glioma network mapping (see Methods), are shown in both volumetric space (left) and surface projections (right) for the Tiantan (*n* = 830), BraTS (*n* = 345) and UPenn (*n* = 135) datasets. In each map, warmer colors (yellow) indicate the cross-individual positive functional overlap of the gliomas, while cooler colors (blue) indicate stronger negative functional overlap of the gliomas. Pearson correlation coefficients comparing the network maps across datasets are shown on the far right (one-sided $P_{spin} < 0.001$).

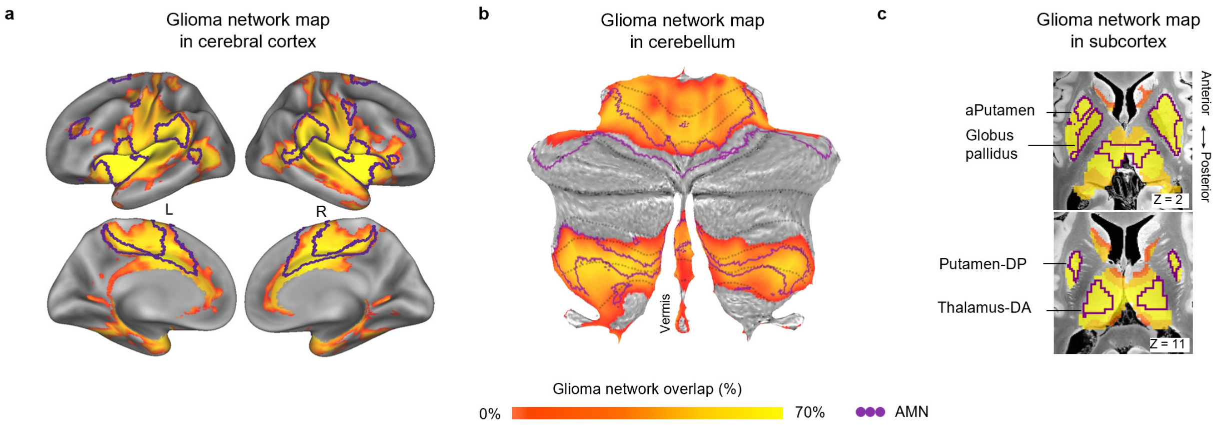


**Supplementary Fig. 2 | Full-threshold display of the cortical and subcortical distribution of the glioma network. a**, Full-threshold glioma network in the cerebral cortex based on all 1,310 patients from the Tiantan, BraTS and UPenn datasets. **b**, Glioma network in the cerebellum (flat map). **c**, Glioma network in subcortical regions. The color scale reflects the overlap of glioma-connected networks across patients, calculated as the percentage of patients exhibiting significant connectivity between the glioma and a given voxel. In a-c, the action-mode network (AMN) is outlined in purple.

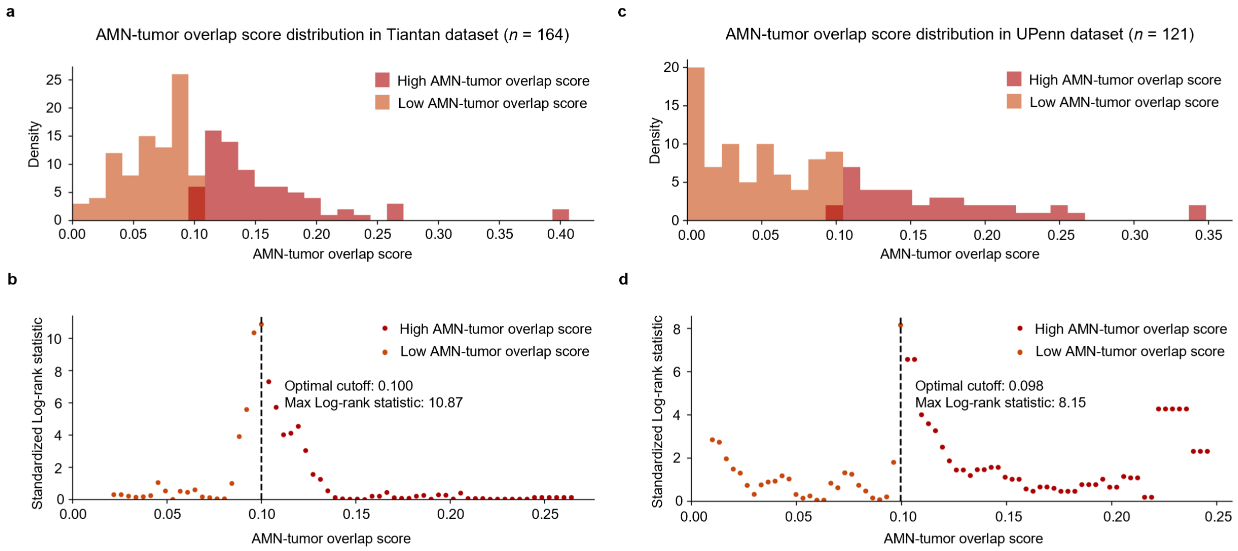


**Supplementary Fig. 3 | Cutoff optimization for action-mode network (AMN)-tumor overlap score.** (a) Histogram showing the distribution of AMN-tumor overlap scores in the Tiantan dataset ($n$ = 164). Patients are stratified into high (red) and low (orange) AMN-tumor overlap score groups. (b) Standardized log-rank statistics corresponding to a range of candidate cutoff values in the Tiantan dataset. The vertical dashed line marks the optimal cutoff (0.100), which yields the maximum log-rank statistic[1] (10.87) for survival separation between the high and low overlap groups. (c) Histogram of AMN-tumor overlap score distribution in the UPenn dataset ($n$ = 121). Patients are stratified into high (red) and low (orange) AMN-tumor overlap score groups. (d) Standardized log-rank statistics corresponding to a range of candidate cutoff values in the UPenn dataset. The optimal cutoff (0.100) yields the maximum log-rank statistic (8.15) for survival separation between the high and low overlap groups.

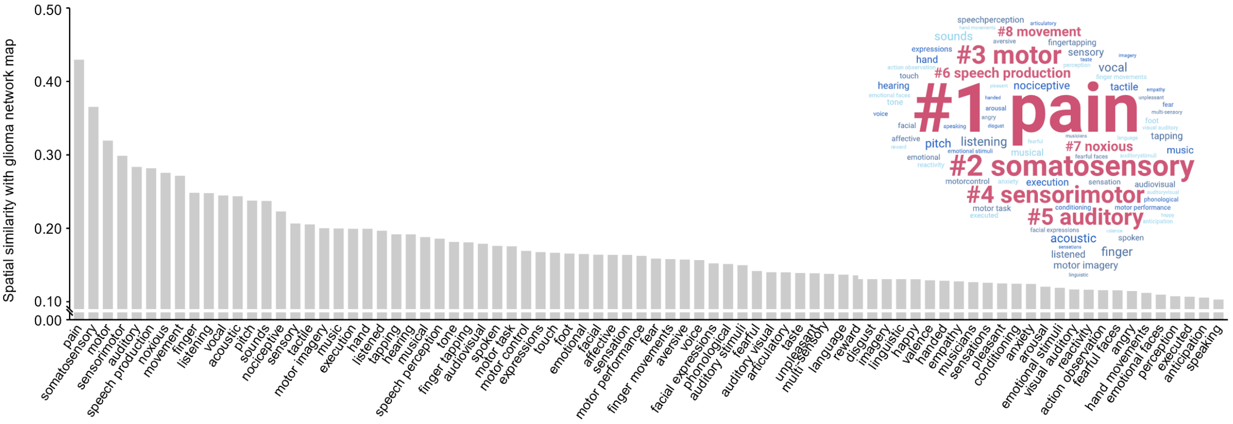


**Supplementary Fig. 4 | Comparative functional pattern analytics based on the NeuroSynth database.** The bar plot (left) shows spatial correlations between the glioma network and behavior-related functional activation maps from the NeuroSynth database[2] (Pearson's $r > 0.100$, $P < 0.001$, FDR-corrected). Each bar represents the strength of spatial similarity for a given functional pattern. The word cloud (right) visualizes functional patterns with Pearson's $r > 0.100$, with word size scaled proportional to correlation strength. The eight functional patterns with the highest similarity to the glioma network are highlighted in bold.

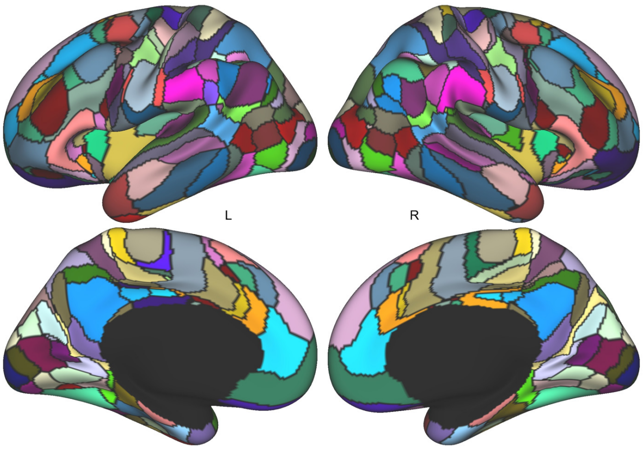


**Supplementary Fig. 5 | Parcellation of 213 brain functional regions.** The cerebral cortex was parcellated into 213 functional regions using a previously described approach[3-5] , with 108 regions in the left hemisphere and 105 regions in the right hemisphere.

### Supplementary Table 1 | Patient demographic and clinical data.

| Characteristic | Tiantan (*n* = 830) | BraTS (*n* = 345) | UPenn (*n* = 135) | Multifocal glioma (*n* = 38) | Cerebellar glioma (*n* = 49) |
|---|---|---|---|---|---|
| Mean age in years (SD) | 44.0 (14.7) | NA | 62.7 (12.0) | 52.4 (14.9) | 43.2 (17.2) |
| Gender, females \| males | 375 \| 455 | NA | 55 \| 80 | 23 \| 15 | 23 \| 26 |
| Glioma localization, left \| right | 388 \| 442 | 181 \| 164 | 67 \| 68 | 16 \| 22 | 18 \| 31 |
| Glioma grade, LGG \| HGG \| NA | 432 \| 398 \| 0 | 74 \| 271 \| 0 | 0 \| 135 \| 0 | 9 \| 24 \| 4 | 28 \| 18 \| 3 |
| Tumor volume in $cm^3$ (SD) | 63 (55) | 53 (45) | 35 (30) | 120 (78) | 27 (22) |
| Molecular subtype, IDH-mut \| IDH-wt \| NA | 477 \| 353 \| 0 | 0 \| 0 \| 345 | 0 \| 109 \| 26 | 3 \| 31 \| 4 | 6 \| 37 \| 6 |
| MGMT, methylated \| unmethylated \| Indeterminate \| NA | 628 \| 143 \| 55 \| 4 | 0 \| 0 \| 0 \| 345 | 21 \| 33 \| 6 \| 75 | 14 \| 12 \| 0 \| 12 | 9 \| 21 \| 0 \| 19 |
| EGFR, amplification \| nonamplification \| NA | 294 \| 8 \| 528 | 0 \| 0 \| 345 | 0 \| 0 \| 135 | 4 \| 13 \| 21 | 1 \| 22 \| 26 |

LGG: low-grade glioma. HGG: high-grade glioma. NA: not available.

IDH: isocitrate dehydrogenase. mut: mutant type. wt: wide type.

MGMT: $O^6$-methylguanine DNA methyltransferase. EGFR: epidermal growth factor receptor.

**Supplementary Table 2 | Specific coordinates of glioma location foci across datasets.** Coordinates represent the top three centroids of regions corresponding to peak glioma occurrence in each dataset. All coordinates are reported in MNI space as [X Y Z].

| Dataset | No. of patients | Coordinates | | | Anatomy of Coordinates |
|---|---|---|---|---|---|
| Tiantan | 120 / 830 (14.5%) | 39 | -9 | -3 | R, long insular gyrus and central insular sulcus |
| | 119 / 830 | 39 | -9 | -5 | R, inferior circular sulcus of the insula |
| | | 39 | -9 | 2 | R, cerebral white matter |
| | 118 / 830 | 47 | -15 | 3 | R, superior temporal gyrus and transverse temporal gyrus |
| BraTS | 45 / 345 (13.0%) | 39 | -25 | 3 | R, cerebral white matter |
| | 44 / 345 | 39 | -33 | 11 | R, cerebral white matter |
| | 43 / 345 | 40 | -25 | -1 | R, cerebral white matter |
| | | -33 | -9 | -7 | L, putamen |
| UPenn | 23 / 135 (17.0%) | 41 | -29 | -2 | R, cerebral white matter |
| | 22 / 135 | 45 | -27 | -5 | R, superior temporal sulcus |
| | 21 / 135 | 38 | -29 | 4 | R, cerebral white matter |

**Supplementary Table 3 | Specific coordinates of glioma network foci across datasets.** Coordinates represent the top three centroids of peak functional connected regions with gliomas in each dataset. Given the broader spatial extent of functional connectivity, a larger interval between patient counts was used. All coordinates are reported in MNI space as [X Y Z].

| Dataset | No. of patients | Coordinates | | | Anatomy of Coordinates |
|---|---|---|---|---|---|
| Tiantan | 634 / 830 (76.4%) | 42 | 6 | -5 | R, short insular gyrus |
| | | 39 | 7 | -5 | R, cerebral white matter |
| | 630 / 830 | -33 | -1 | -2 | L, putamen |
| | 625 / 830 | 39 | -9 | -5 | R, inferior circular sulcus of the insula |
| BraTS | 242 / 345 (70.1%) | 39 | -1 | -4 | R, long insular gyrus and central insular sulcus |
| | 238 / 345 | 41 | -5 | -4 | R, inferior circular sulcus of the insula |
| | 235 / 345 | 39 | -12 | -4 | R, inferior circular sulcus of the insula |
| | | 41 | 0 | -4 | R, short insular gyrus |
| UPenn | 96 / 135 (71.1%) | 39 | -1 | -13 | R, inferior circular sulcus of the insula |
| | 94 / 135 | -34 | -1 | -12 | L, putamen |
| | | 46 | -16 | -4 | R, cerebral white matter |
| | 92 / 135 | 39 | 1 | -13 | R, long insular gyrus and central insular sulcus |

**Supplementary Table 4 | Specific coordinates of overall glioma network foci.** Coordinates represent the top three centroids of peak functional connected regions with gliomas in each dataset. All coordinates are reported in MNI space as [X Y Z].

| | **Anatomy of Coordinates** | | **No. of patients** | **Coordinates** | | |
|---|---|---|---|---|---|---|
| Cerebral Cortex | Putamen | L | 944 / 1310 (72.1%) | -33 | -17 | 3 |
| | inferior circular sulcus of the insula | R | 952 / 1310 (72.7%) | 39 | -9 | -5 |
| Cerebellum | VIIA / VIIB | L | 816 / 1310 (62.3%) | -17 | -65 | -45 |
| | | R | 801 / 1310 (61.1%) | 23 | -65 | -45 |
| | V / VI | L | 763 / 1310 (58.2%) | -17 | -58 | -21 |
| | | R | 742 / 1310 (56.6%) | 23 | -57 | -21 |
| Subcortex | Putamen | L | 930 / 1310 (71.0%) | -31 | -1 | 3 |
| | | R | 910 / 1310 (69.5%) | 34 | -1 | -4 |
| | Globus pallidus | L | 911 / 1310 (69.6%) | -24 | -9 | -5 |
| | | R | 882 / 1310 (67.4%) | 23 | -1 | -5 |
| | Thalamas DA | L | 853 / 1310 (65.1%) | -9 | -17 | 3 |
| | | R | 874 / 1310 (66.7%) | 15 | -17 | 3 |

DA: dorsoanterior.

**Supplementary Table 5 | Glioma risk scores across canonical functional networks.** Glioma risk score is defined as cross-participant overlap of glioma-connected networks (positive value) and their anti-correlations (negative value). Glioma risk scores for each canonical network were computed separately across different brain regions (whole brain, cerebral cortex, cerebellum, and subcortex) based on all 1,310 patients from the Tiantan, BraTS and UPenn datasets. Subgroup analyses were performed using the Tiantan dataset for specific tumor types, including low-grade gliomas (LGG, $n$ = 432), high-grade gliomas (HGG, $n$ = 398), IDH-mutant gliomas (IDH-mut, $n$ = 477), and IDH-wild gliomas (IDH-wt, $n$ = 353). Values are presented as mean (standard deviation).

| | **AMN** | **SMN** | **LMB** | **dATN** | **DMN** | **FPN** | **VIS** |
|---|---|---|---|---|---|---|---|
| Whole brain | 0.445 (0.276) | 0.299 (0.385) | -0.062 (0.413) | -0.179 (0.410) | -0.256 (0.409) | -0.339 (0.366) | -0.493 (0.354) |
| Cerebral cortex | 0.508 (0.230) | 0.309 (0.390) | -0.138 (0.403) | -0.155 (0.390) | -0.235 (0.401) | -0.284 (0.395) | -0.438 (0.353) |
| Cerebellum | 0.230 (0.295) | 0.337 (0.218) | 0.185 (0.302) | 0.105 (0.289) | -0.463 (0.243) | -0.371 (0.289) | 0.005 (0.274) |
| Subcortex | 0.615 (0.046) | 0.614 (0.068) | 0.429 (0.236) | 0.560 (0.046) | 0.379 (0.247) | 0.399 (0.258) | 0.521 (0.032) |
| LGG | 0.572 (0.178) | 0.222 (0.450) | -0.115 (0.407) | -0.289 (0.403) | -0.158 (0.428) | -0.137 (0.467) | -0.608 (0.224) |
| HGG | 0.470 (0.298) | 0.294 (0.395) | -0.186 (0.419) | -0.107 (0.391) | -0.318 (0.377) | -0.362 (0.370) | -0.238 (0.459) |
| IDH-mut | 0.599 (0.155) | 0.285 (0.423) | -0.228 (0.362) | -0.216 (0.427) | -0.220 (0.413) | -0.128 (0.474) | -0.605 (0.223) |
| IDH-wt | 0.403 (0.340) | 0.187 (0.430) | -0.088 (0.440) | -0.216 (0.370) | -0.247 (0.416) | -0.383 (0.361) | -0.227 (0.467) |

LGG: low-grade glioma. HGG: high-grade glioma.

IDH: isocitrate dehydrogenase. mut: mutant type. wt: wide type.

**Supplementary Table 6 | Demographics and clinical summary of patient datasets used for survival analyses.**

Tiantan Dataset ($n$ = 164)

| | High-overlap | Low-overlap |
|---|---|---|
| No. of patients | 75 | 89 |
| Mean age in years (SD) | 46.0 (13.6) | 44.9 (12.7) |
| Gender, females \| males | 38 \| 37 | 35 \| 54 |
| Glioma grade (LGG \| HGG) | 41 \| 34 | 52 \| 37 |
| Tumor volume in $cm^3$ (SD) | 96.9 (83.8) | 90.6 (85.0) |

LGG: low-grade glioma. HGG: high-grade glioma.

UPenn Dataset ($n$ = 121)

| | High-overlap | Low-overlap |
|---|---|---|
| No. of patients | 43 | 78 |
| Mean age in years (SD) | 61.6 (9.20) | 59.9 (11.1) |
| Gender, females \| males | 18 \| 25 | 33 \| 45 |
| Tumor volume in $cm^3$ (SD) | 39.3 (34.1) | 31.0 (27.5) |

**Supplementary Table 7 | Univariate and multivariate survival analysis of patient datasets.**

| Characteristic | Univariate Cox | | Multivariate Cox | |
|---|---|---|---|---|
| | HR (95% CI) | *P*-value | HR (95% CI) | *P*-value |
| Age | 1.022 (1.006, 1.037) | **0.0061** | 1.018 (1.001, 1.034) | **0.033** |
| Gender = female | 1.037 (0.702, 1.531) | 0.856 | 1.018 (0.684, 1.515) | 0.930 |
| Glioma Grade = HGG | 1.199 (0.988, 1.456) | 0.066 | 1.083 (0.873, 1.343) | 0.469 |
| Tumor Volume | 1.001 (0.999, 1.003) | 0.339 | 1.001 (0.999, 1.003) | 0.373 |
| AMN-tumor overlap score = high | 1.906 (1.290, 2.815) | **0.0012** | 1.829 (1.236, 2.707) | **0.0025** |

HGG: high-grade glioma.

Cox proportional hazards model was performed for univariate and multivariate regression (Tiantan *n* = 164).

| Characteristic | Univariate Cox | | Multivariate Cox | |
|---|---|---|---|---|
| | HR (95% CI) | *P*-value | HR (95% CI) | *P*-value |
| Age | 1.022 (1.005, 1.039) | **0.010** | 1.025 (1.007, 1.043) | **0.0057** |
| Gender = female | 1.180 (0.820, 1.698) | 0.373 | 1.305 (0.901, 1.892) | 0.159 |
| Tumor Volume | 0.999 (0.994, 1.005) | 0.745 | 1.000 (0.994, 1.005) | 0.903 |
| AMN-tumor overlap score = high | 1.803 (1.227, 2.650) | **0.0027** | 1.825 (1.239, 2.688) | **0.0023** |

Cox proportional hazards model was performed for univariate and multivariate regression (UPenn *n* = 121).

**Supplementary Table 8 | Details of neurotransmitters and other metabolism measures datasets.**

| Feature | Description | Primary Reference |
|---|---|---|
| VAChT | PET tracer binding (SUVR) to VAChT (acetylcholine transporter) | Aghourian et al., 2017 [6] |
| 5-HTT | PET tracer binding (Bmax) to 5-HTT (serotonin transporter) | Beliveau et al., 2017 [7] |
| DAT | SPECT tracer binding (SUVR) to DAT (dopamine transporter) | Dukart et al., 2018 [8] |
| 5-$HT_{1A}$ | PET tracer binding (Bmax) to 5-$HT_{1A}$ (serotonin receptor) | Beliveau et al., 2017 [7] |
| $D_2$ | PET tracer binding (BPnd) to $D_2$ (dopamine receptor) | Jaworska et al., 2020 [9]<br>Hansen et al., 2021 [10] |
| $H_3$ | PET tracer binding (Vt) to $H_3$ (histamine receptor) | Gallezot et al., 2017 [11]<br>Hansen et al., 2021 [10] |
| MOR | PET tracer binding (BPnd) to MOR (mu-opioid receptor) | Kantonen et al., 2020 [12] |
| $CB_1$ | PET tracer binding (Vt) to $CB_1$ (cannabinoid receptor) | Normandin et al., 2015 [13] |
| 5-$HT_4$ | PET tracer binding (Bmax) to 5-$HT_4$ (serotonin receptor) | Beliveau et al., 2017 [7] |
| $D_1$ | PET tracer binding (BPnd) to $D_1$ (dopamine receptor) | Kaller et al., 2017 [14] |
| NET | PET tracer binding (BPnd) to NET (norepinephrine transporter) | Ding et al., 2010 [15]<br>Hansen et al., 2021 [10] |
| $\alpha_4\beta_2$ | PET tracer binding (Vt) to $\alpha_4\beta_2$ (acetylcholine receptor) | Hillmer et al., 2016 [16]<br>Hansen et al., 2021 [10] |
| $mGluR_5$ | PET tracer binding (BPnd) to $mGluR_5$ (glutamate receptor) | Hansen et al., 2021 [10] |
| 5-$HT_6$ | PET tracer binding (BPnd) to 5-HT6 (serotonin receptor) | Radhakrishnan et al., 2018 [17] |
| 5-$HT_{2A}$ | PET tracer binding (Bmax) to 5-$HT_{2A}$ (serotonin receptor) | Beliveau et al., 2017 [7] |
| $M_1$ | PET tracer binding (BPnd) to $M_1$ (acetylcholine receptor) | Naganawa et al., 2020 [18]<br>Hansen et al., 2021 [10] |

*(continued on next page)*

| Feature | Description | Primary Reference |
| --- | --- | --- |
| 5-$HT_{1B}$ | PET tracer binding (Bmax) to 5-$HT_{1B}$ (serotonin receptor) | Beliveau et al., 2017 [7] |
| $GABA_A$ | PET and autoradiography informed $GABA_A$ benzodiazepine binding-site density (Bmax; GABA receptor) | Norgaard et al., 2021 [19] |
